# An integrated RNA and CRISPR/Cas toolkit for multiplexed synthetic circuits and endogenous gene regulation in human cells

**DOI:** 10.1101/004432

**Authors:** Lior Nissim, Samuel D. Perli, Alexandra Fridkin, Pablo Perez-Pinera, Timothy K. Lu

## Abstract

RNA-based regulation, such as RNA interference, and CRISPR/Cas transcription factors (CRISPR-TFs), can enable scalable synthetic gene circuits and the modulation of endogenous networks but have yet to be integrated together. Here, we combined multiple mammalian RNA regulatory strategies, including RNA triple helix structures, introns, microRNAs, and ribozymes, with Cas9-based CRISPR-TFs and Cas6/Csy4-based RNA processing in human cells. We describe three complementary strategies for expressing functional gRNAs from transcripts generated by RNA polymerase II (RNAP II) promoters while allowing the harboring gene to be translated. These architectures enable the multiplexed expression of proteins and multiple gRNAs from a single compact transcript for efficient modulation of synthetic constructs and endogenous human promoters. We used these regulatory tools to implement tunable synthetic gene circuits, including multi-stage transcriptional cascades. Finally, we show that Csy4 can rewire regulatory connections in RNA-dependent gene circuits with multiple outputs and feedback loops to achieve complex functional behaviors. This multiplexable toolkit will be valuable for the construction of scalable gene circuits and the perturbation of natural regulatory networks in human cells for basic biology, therapeutic, and synthetic-biology applications.

## Introduction

The ability to build complex, robust, and scalable synthetic gene networks that operate with defined interconnections between artificial parts and native cellular processes is central to engineering biological systems. Furthermore, this capability can enable new strategies for rewiring, perturbing, and probing natural biological networks. A large set of tunable, orthogonal, compact, and multiplexable gene regulatory mechanisms is of fundamental importance to implement these applications. However, despite much progress in the fields of transcriptional regulation and synthetic biology, the tools that are available at this time fail to meet one or more of the criteria described above. For example, transcriptional regulation utilizes transcription factors that bind predetermined DNA sequences of interest. Previously, natural DNA-binding proteins have been used to target effector domains, such as activators and repressors, domains to the regulatory regions of mammalian genes to modulate their transcription (Cronin et al., 2001; Gossen and Bujard, 1992). However, only a few orthogonal variants of natural DNA-binding proteins are characterized well and modifying their sequence specificity is challenging (Urlinger et al., 2000). Other approaches have focused on engineering artificial DNA-binding proteins such as zinc fingers (ZFs) and transcription-activator-like effectors (TALEs), which can require multi-step assembly, screening, and/or optimization processes to achieve desired DNA binding characteristics (Beerli and Barbas, 2002; Blount et al., 2012; Khalil et al., 2012; Lohmueller et al., 2012; Maeder et al., 2009; Reyon et al., 2012; Sanjana et al., 2012).

Recently, type II CRISPR/Cas systems (DNA-targeting Cas proteins) have been adapted to achieve programmable DNA binding without requiring complex protein engineering (Sander and Joung, 2014). In these systems, the sequence specificity of the Cas9 DNA-binding protein is determined by guide RNAs (gRNAs) which have base-pairing complementarity to target DNA sites. This enables simple and highly flexible programing of Cas9 binding. Cas9’s nuclease activity has been utilized for precise and efficient genome editing in prokaryotic and eukaryotic cells (Cong et al., 2013; Jiang et al., 2013; Jinek et al., 2012; Jinek et al., 2013; Mali et al., 2013b; Wang et al., 2013). More recently, a mutant derivative of this protein (dCas9), which lacks nuclease activity, was modified to enable programmable transcriptional regulation of both ectopic and native promoters to create CRISPR-based transcription factors (CRISPR-TFs) in mammalian cells (Cheng et al., 2013; Farzadfard et al., 2013; Gilbert et al., 2013; Maeder et al., 2013a; Mali et al., 2013a; Perez-Pinera et al., 2013a). Type III CRISPR/Cas systems (RNA-targeting Cas proteins) have also been adapted for synthetic-biology applications (Qi et al., 2012). For example, the type III CRISPR/Cas-associated Csy4 protein from *Pseudomonas aeruginosa* has been used in bacteria to achieve predictable regulation of multi-gene operons by cleaving precursor mRNAs (Qi et al., 2012). The functionality of the Csy4 protein has also been demonstrated in bacteria, archaea, and eukaryotes (Qi et al., 2012).

CRISPR-TFs have the potential to enable the scalable construction of large-scale synthetic gene circuits and the rewiring of natural regulatory networks. This is due to the ease by which one can define new, orthogonal transcriptional regulators by designing artificial gRNAs. However, up until now, gRNAs for gene regulation in human cells have only been expressed from RNA polymerase III (RNAP III) promoters. This is an important limitation in terms of integrating CRISPR/Cas regulation with endogenous gene networks. This is because RNAP III promoters comprise only a small portion of cellular promoters and are mostly constitutively active (Orioli et al., 2012; Teichmann et al., 2010; White, 1998; Willis, 1993), thus preventing the linkage of most cellular promoters and signals into CRISPR-TF-based networks. Furthermore, multiple gRNAs are typically needed to efficiently activate endogenous promoters (Cheng et al., 2013; Maeder et al., 2013a; Mali et al., 2013a; Perez-Pinera et al., 2013a) but strategies for multiplexed gRNA production from single transcripts for transcriptional regulation have not yet been described. As a result, multiple gRNA expression constructs are currently needed to perturb natural transcriptional networks, thus limiting scalability. In addition to transcriptional regulation, natural circuits leverage RNA-based translational and post-translational regulation to achieve complex behavior (Audibert et al., 2002; Chen and Manley, 2009; Feng et al., 2006; Lee, 2012; Lin et al., 2006; Mercer et al., 2009; Nagano et al., 2008; Pandey et al., 2008; Rinn et al., 2007; Saito et al., 2011; Wang et al., 2008; Wilson and Doudna, 2013; Zhao et al., 2008). Thus, synthetic gene regulatory strategies that combine RNA and transcriptional engineering could be useful in modeling natural systems or implementing artificial behaviors.

Here, we present a flexible toolkit that integrates mammalian and bacterial RNA-based regulatory mechanisms to create complex synthetic circuit topologies and to regulate endogenous promoters. We used multiple mammalian RNA processing strategies, including 3’ RNA triple helixes (triplexes) (Wilusz et al., 2012), introns, and ribozymes, together with mammalian miRNA regulation, bacteria-derived CRISPR-TFs, and the Csy4 RNA-modifying protein from *P. aeruginosa*. We used these tools to generate functional gRNAs from RNAP-II-regulated mRNAs in human cells while rendering the concomitant translation of the harboring mRNAs tunable. These functional gRNAs were used to target both synthetic and endogenous promoters for activation via CRISPR-TFs. Furthermore, we developed strategies for multiplexed gRNA production, thus enabling compact encoding of proteins and multiple gRNAs in single transcripts. To demonstrate the utility of these regulatory parts, we implemented multi-stage transcriptional cascades that are necessary for the construction of complex synthetic gene circuits. We also combined mammalian miRNA-based regulation with CRISPR-TFs to create multi-component genetic circuits whose feedback loops, interconnections, and behaviors could be rewired by Csy4-based RNA processing. Thus, this platform can be used to construct, synchronize, and switch complex regulatory networks, both artificial and endogenous, using synthetic transcriptional and RNA-dependent mechanisms. We envision that the integration of CRISPR-TF-based gene regulation systems with mammalian RNA regulatory architectures should enable scalable gene regulatory systems for synthetic biology as well as basic biology applications.

## Results

### Functional gRNA generation with an RNA triple helix and Csy4

An important first step to enabling complex CRISPR-TF-based circuits is to generate functional gRNAs from RNAP II promoters in human cells, which would allow for the coupling of gRNA production to specific regulatory signals. For example, the activation of gRNA-dependent circuits could be initiated in defined cell types or states, or in response to external inputs. Furthermore, the ability to simultaneously express gRNAs along with proteins from a single transcript would be beneficial. This would enable multiple outputs, including effector proteins and regulatory links, to be produced from a concise genetic architecture. It could also enable the integration of gRNA expression into endogenous loci. Thus, we sought to create a system in which functional gRNAs and proteins could be simultaneously produced by endogenous RNAP II promoters.

We utilized the RNA-binding and RNA-endonuclease capabilities of the Csy4 protein from *P. aeruginosa* (Haurwitz et al., 2012; Sternberg et al., 2012). Csy4 recognizes a 28 nucleotide RNA sequence (hereafter referred to as the ‘28’ sequence), cleaves the RNA, and remains bound to the upstream RNA fragment (Haurwitz et al., 2012). We thus utilized Csy4 to release gRNAs from transcripts generated by RNAP II promoters which also encode functional protein sequences. To generate a gRNA-harboring transcript, we used the potent CMV promoter (CMVp) to express the mKate2 protein. We encoded a gRNA (gRNA1), flanked by two Csy4 binding sites, downstream of the coding region of mKate2 (Figure 1A). In this architecture, RNA cleavage by Csy4 releases a functional gRNA but also removes the poly-(A) tail from the upstream mRNA (encoding mKate2 in this case), which is known to result in impaired translation of most eukaryotic mRNAs (Jackson, 1993; Proudfoot, 2011).

**Figure 1.**
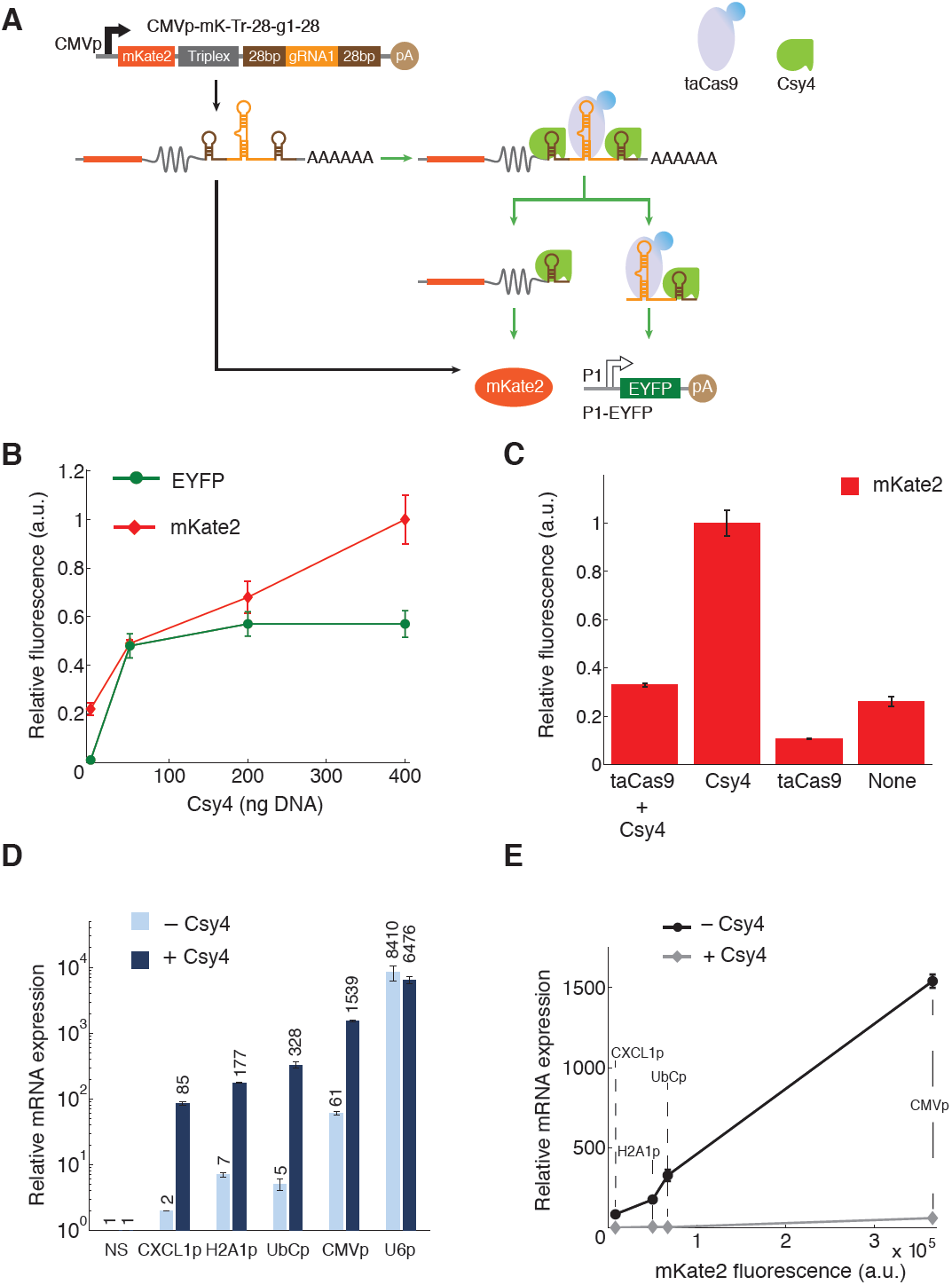
The ‘triplex/Csy4’ architecture (CMVp-mK-Tr-28-g1-28) produces functional gRNAs from RNAP II promoters while maintaining expression of the harboring gene. The gRNAs can activate synthetic promoters (A-C) as well as endogenous promoters (D-E). **(A)** gRNA1 was flanked by two Csy4 recognition sites (‘28’) and placed down-stream of an mKate2 gene followed by an RNA triple helix (triplex). The entire transcript was expressed by a CMV promoter (CMVp). Csy4 enables the generation of functional gRNAs which can be incorporated into a transcriptionally active dCas9-VP64 (taCas9) that can activate a synthetic promoter (P1) driving EYFP expression (P1-EYFP). **(B)** The presence of Csy4 enabled a 60-fold increase in EYFP levels, demonstrating the generation of functional gRNAs. Furthermore, increased concentrations of the Csy4-expressing plasmid led to increasing mKate2 levels. Fluorescence values were normalized to the maximum respective fluorescence between the data in this figure and Figure 2B-D to enable cross comparisons between the ‘triplex/Csy4’ and ‘intron/Csy4’ architectures. **(C)** Csy4 and taCas9 have opposite effects on mKate2 fluorescence generated by the CMVp-mK-Tr-28-g1-28 construct. The taCas9 construct alone reduced mKate2 levels while the Csy4 construct alone enhanced mKate2 fluorescence. The mKate2 expression levels were normalized to the maximum mKate2 value observed (Csy4 only) across the four conditions tested here. **(D)** The human RNAP II promoters, CXCL1p, H2A1p, and UbCp, as well as the RNAP II promoter CMVp were used to drive expression of four different gRNAs (gRNA3-6, Table S2) previously shown to activate the IL1RN promoter (Perez-Pinera et al., 2013a) from the ‘triplex/Csy4’ construct. These results were compared to the RNAP III promoter U6p driving direct expression of the same gRNAs. Four different plasmids, each containing one of the indicated promoters and gRNAs 3-6, were co-transfected along with a plasmid encoding taCas9 and with or without a plasmid expressing Csy4. Relative IL1RN mRNA expression, compared to a control construct with non-specific gRNA (NS, CMVp-mK-Tr-28-g1-28), was monitored using qRT-PCR. The RNAP II promoters resulted in a wide range of IL1RN activation, with the presence of Csy4 greatly increasing activation compared with the absence of Csy4. Interestingly, IL1RN activation was achieved by the RNAP II promoters even in the absence of Csy4, albeit at much lower levels than in the presence of Csy4. **(E)** The input-output transfer curve for the activation of the endogenous IL1RN loci by the ‘triplex/Csy4’ construct was determined by plotting the mKate2 levels (as a proxy for the input) versus the relative IL1RN mRNA expression levels (as the output). The data indicate that tunable modulation of endogenous loci can be achieved with RNAP II promoters of different strengths. The IL1RN data is the same as shown in D).

To enable efficient translation of mRNA lacking a poly-(A) tail, we sought to identify RNA components capable of functionally complementing the loss of the poly-(A). We cloned a 110 bp fragment derived from the 3’ end of the mouse MALAT1 locus (Wilusz et al., 2012) downstream of mKate2 and immediately upstream of the gRNA sequence flanked by Csy4 recognition sites. The MALAT1 lncRNA is deregulated in many human cancers (Lin et al., 2006) and despite lacking a poly-(A) tail, the MALAT1 is a stable transcript (Wilusz et al., 2008; Wilusz et al., 2012) which is protected from the exosome and 3’-5’ exonucleases by a highly conserved 3’ triple helical structure (triplex) (Wilusz et al., 2012). Indeed, a 3’ triplex sequence has been shown to be sufficient for *in vivo* translation of transcripts lacking a poly-(A) tail (Wilusz et al., 2012). Thus, our final ‘triplex/Csy4’ architecture was a CMVp-driven mKate2 transcript with a 3’ triplex sequence followed by a 28-gRNA-28 sequence (CMVp-mK-Tr-28-gRNA-28) (Figure 1A).

To characterize gRNA activity, we co-transfected HEK-293T cells with the CMVp-mK-Tr-28-gRNA1-28 expression plasmid, along with a plasmid encoding a synthetic P1 promoter that is specifically activated by gRNA1 to express EYFP. The P1 promoter contains 8x binding sites for gRNA1 and is based on a minimal promoter construct (Farzadfard et al., 2013). In this experiment and all the following ones (unless otherwise indicated), we also co-transfected a transcriptionally active dCas9-NLS-VP64 protein (taCas9) expressed by a CMV promoter. We transfected HEK-293T cells with 0-400 ng of a Csy4-expressing plasmid (where Csy4 was produced by the murine PGK1 promoter) along with 1 µg of the other plasmids (Figure 1B and Figure S1A for raw data).

Increasing Csy4 concentration levels did not result in a decrease of mKate2 levels, but instead led to an up to 5-fold increase (Figure 1B). Furthermore, functional gRNAs generated from this construct induced EYFP expression by up to 60-fold from the P1 promoter. While mKate2 expression continued to increase with the concentration of the Csy4-expressing plasmid, EYFP activation plateaued after 50 ng of the Csy4-producing plasmid. In addition, we observed evidence of cytotoxicity at 400 ng Csy4 plasmid concentrations. Thus, we decided to use 100-200 ng of the Csy4 plasmid in subsequent experiments (except where otherwise noted), although this reduced the number of Csy4-positive cells after transfection. This issue could be addressed in future work by using weaker promoters to reduce Csy4 expression levels or by generating stable cell lines with low or moderate levels of Csy4.

Interestingly, although a 5’ Csy4 recognition site alone should be sufficient to release gRNAs from the harboring RNA transcript, this variant architecture did not generate functional gRNAs capable of activating a downstream target promoter above background levels (data not shown). This could be the result of RNA destabilization, poly-(A)-mediated cytoplasmic transport, interference of the poly-(A) tail with taCas9 activity, or other yet to be determined mechanisms.

We further characterized the relative effects of Csy4 and taCas9 on the expression of mKate2. We measured mKate2 fluorescence from the ‘triplex/Csy4’-based gRNA expression construct in the presence of Csy4 and taCas9, Csy4 alone, taCas9 alone, or neither protein (Figure 1C and Figure S2). The lowest mKate2 fluorescence levels resulted from the taCas9 only condition. Since we used a taCas9 with a strong nuclear localization sequence (NLS), this effect could arise from taCas9 binding to the gRNA within the mRNA and localizing the transcript to the nucleus. This hypothesis is supported by data demonstrating that endogenous promoters can be activated by gRNAs produced from the ‘triplex/Csy4’-based architecture even in the absence of Csy4 (see below and Figure 1D,E). The highest mKate2 expression levels were obtained with Csy4 alone, suggesting that Csy4 processing could enhance mKate2 levels. Expression of mKate2 in the absence of both Csy4 and taCas9 as well as in the presence of both Csy4 and taCas9 were similar and reduced by 3-4 fold compared with Csy4 only.

### Modulating endogenous loci with CRISPR-TFs expressed from human promoters

To validate the robustness of the ‘triplex/Csy4’ architecture, we adapted it to regulate the expression of a native genomic target in human cells. It has been previously shown that combinations of synthetic transcription factors recruited to endogenous human promoters lead to synergistic and robust activation of gene expression (Cheng et al., 2013; Maeder et al., 2013a; Maeder et al., 2013b; Mali et al., 2013a; Perez-Pinera et al., 2013a; Perez-Pinera et al., 2013b). Hence, we targeted the endogenous IL1RN locus for gene activation via the co-expression of four distinct gRNAs, gRNA3-6 (Table S2) (Perez-Pinera et al., 2013a).

We designed each of the four gRNAs to be expressed concomitantly with mKate2, each from a separate plasmid. Each set of four gRNAs was regulated by one of the following promoters (in descending order according to their activity level in HEK-293T cells): the Cytomegalovirus Immediate Early (CMVp), human Ubiquitin C (UbCp), human Histone H2A1 (H2A1p) (Rogakou et al., 1998), and human inflammatory chemokine CXCL1 (CXCL1p) promoters (Wang et al., 2006). As a control, we also used the RNAP III promoter U6 (U6p) to drive expression of the four gRNAs. For each promoter tested, four plasmids encoding the four different gRNAs were co-transfected along with plasmids expressing taCas9 and Csy4. As a negative control, we substituted the IL1RN-targeting gRNA expression plasmids with plasmids that expressed gRNA1 which was non-specific for the IL1RN promoter (Figure 1D, ‘NS’).

We used qRT-PCR to quantify the mRNA levels of the endogenous IL1RN gene, with the results normalized to the negative control. With the four gRNAs regulated by the U6 promoter, IL1RN activation levels were increased by 8,410-fold in the absence of Csy4 and 6,476-fold with 100 ng of the Csy4-expressing plasmid over the negative control (Figure 1D, ‘U6p’). IL1RN activation with gRNAs expressed from the CMV promoter was substantial (Figure 1D, ‘CMVp’), with 61-fold enhancement in the absence of Csy4 and 1539-fold enhancement with Csy4. The human RNAP II promoters generated ∼2-7 fold activation in the absence of Csy4 and ∼85-328-fold activation with Csy4 (Figure 1D, ‘CXCL1p’, ‘H2A1p’, ‘UbCp’).

To further characterize the input-output transfer function of endogenous gene regulation, we used mKate2 fluorescence generated by each promoter as a marker of input promoter activity for the various RNAP II promoters (Figure 1E). The resulting transfer function was nearly linear in IL1RN activation over the range of mKate2 tested. This data indicates that IL1RN activation was not saturated in the conditions tested and that a large dynamic range of endogenous gene regulation can be achieved with human RNAP II promoters. Thus, tunable modulation of native genes can be achieved using CRISPR-TFs with gRNAs expressed from the ‘triplex/Csy4’ architecture.

### Functional gRNA generation from introns with Csy4

As a complement to the ‘triplex/Csy4’ architecture, we developed an alternative strategy for generating functional gRNAs from RNAP II promoters by encoding a gRNA within an intron in the coding sequence of a gene. This strategy has been used generate synthetic siRNAs in mammalian cells (Greber et al., 2008). Specifically, we encoded gRNA1 as an intron within the coding sequence of mKate2 (Figure 2A) using previously described ‘consensus’ acceptor, donor, and branching sequences (Smith et al., 1989; Taggart et al., 2012). We expected that, once spliced, the gRNA would associate with taCas9 to activate a synthetic promoter P1 regulating EYFP. However, this simple architecture resulted in undetectable EYFP levels (Figure S3, bottom panel). This data is consistent with previous studies highlighting that the lifetime of most intronic RNAs is typically only a few minutes (Audibert et al., 2002; Clement et al., 1999), although the native cellular machinery can generate functional RNA for RNAi from introns (Kim and Kim, 2007; Ying and Lin, 2005). Thus, we concluded that without any stabilization, intronic gRNAs would be expected to be rapidly degraded.

**Figure 2.**
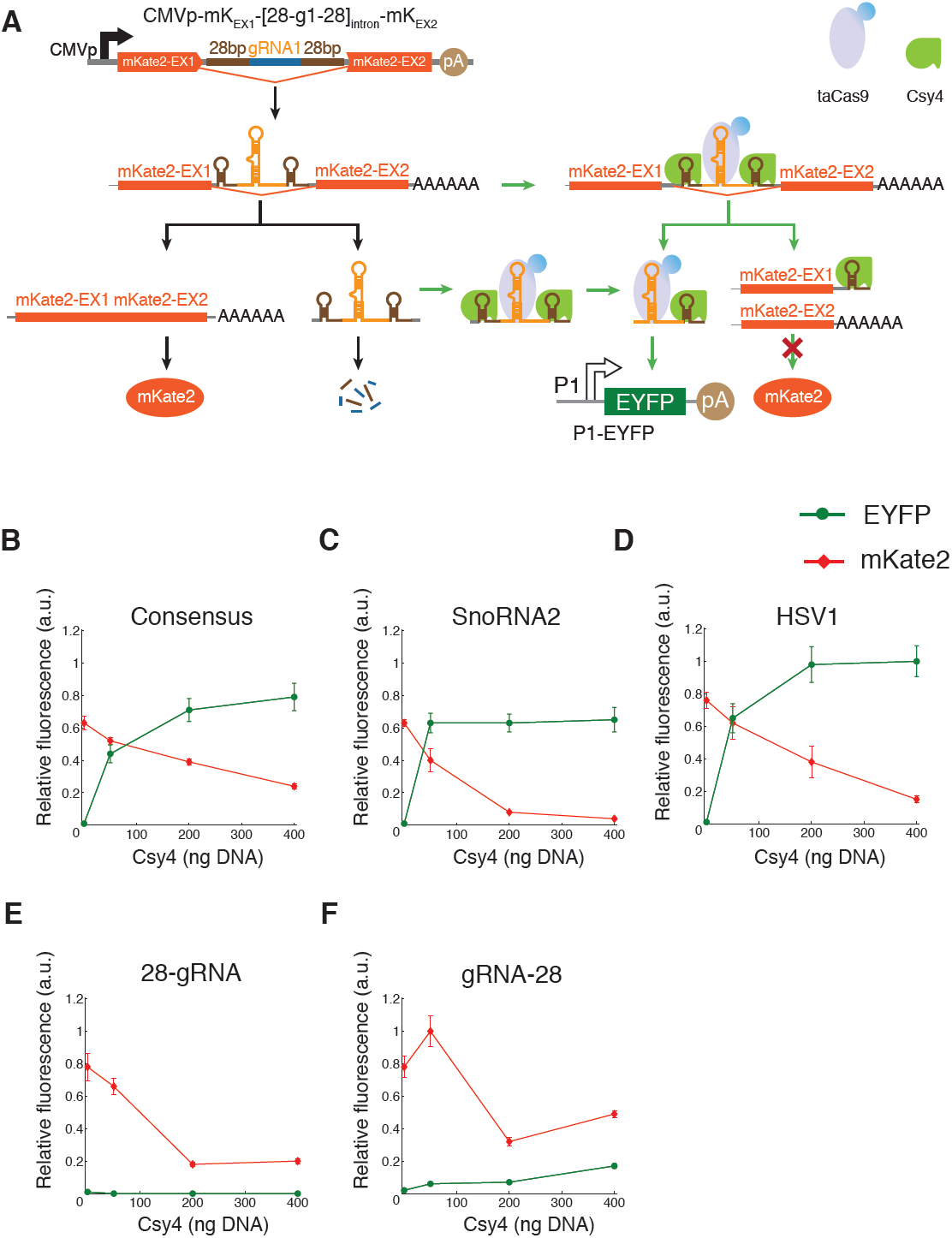
The ‘intron/Csy4’ architecture (CMVp-mK_EX1_-[28-g1-28]_intron_-mK_EX2_) generates functional gRNAs from introns expressed by RNAP-II-expressed transcripts while maintaining modulation of the harboring gene. **(A)** gRNA1 is flanked by Csy4 recognition sites and encoded within an intron, leading to functional gRNA1 generation with Csy4 and activation of a downstream P1-EYFP construct. In contrast to the ‘triplex/Csy4’ construct in Figure 1, the ‘intron/Csy4’ architecture leads to decreased expression of the harboring gene with increased Csy4 levels, which may be due to cleavage of pre-mRNA prior to splicing. **(B-D)** Three introns, a consensus intron (B), snoRNA2 intron (C), and an HSV1 intron (D), combined with Csy4, resulted in functional gRNAs as assessed by EYFP expression. Fluorescence values were normalized to the maximum fluorescence between this data and Figure 1B. **(E)** A single Csy4 binding site located upstream of the gRNA within an HSV1 intron did not produce functional gRNAs but did lead to reduced mKate2 fluorescence with greater Csy4 levels. **(F)** A single Csy4 binding site located downstream of the gRNA within an HSV1 intron produced low levels of functional gRNA and also generated reduced mKate2 levels with greater Csy4-expressing plasmid concentrations. The fluorescence values for (E-F) were normalized to the maximum fluorescence levels between these experiments and a [28-g1-28]_HSV1_ control (Figure S4).

We tried two different approaches to stabilize intronic gRNAs. First, we used intronic sequences that have been reported to produce long-lived introns. This included constructs such as the HSV-1 latency associated intron, which forms a stable circular intron (Block and Hill, 1997), and the sno-lncRNA2 (snoRNA2) intron. The snoRNA2 intron is processed on both ends by the snoRNA machinery, which protects it from degradation and leads to the accumulation of lncRNAs flanked by snoRNA sequences which lack 5′ caps and 3′ poly-(A) tails. (Yin et al., 2012). However, these approaches for generating stable intronic gRNAs also resulted in undetectable activation of the target promoter (data not shown).

As an alternative strategy, we then sought to stabilize intronic gRNAs by flanking the gRNA cassette with two Csy4 recognition sites. In this model, spliced gRNA-containing introns should be bound by Csy4, which should release functional gRNAs. In contrast to the ‘triplex/Csy4’ setting, Csy4 can also potentially bind and digest the pre-mRNA before splicing occurs. In this case, functional gRNA would be produced, but the mKate-containing pre-mRNA would be destroyed in the process (Figure 2A). Thus, increased Csy4 concentrations would be expected to result in decreased mKate2 levels but greater levels of functional gRNA. In this architecture, the decrease in mKate2 levels and increase in functional gRNA with Csy4 concentrations could be expected to depend on several factors, which are illustrated in Figure 2A (black lines, Csy4-independent processes; green lines, Csy4-mediated processes). These competing factors include the rate at which Csy4 binds to its target sites and cleaves the RNA, the rate of splicing, and the rate of spliced gRNA degradation in the absence of Csy4. To examine the behavior of the ‘intron/Csy4’ architecture, we used the CMV promoter to drive expression of mKate2 harboring HSV1, snoRNA, and consensus introns containing gRNA1 flanked by two Csy4-binding-sites (CMVp-mK_EX1_-[28-g1-28]_intron_-mK_EX2_) along with a synthetic P1 promoter regulating the expression of EYFP (Figure 2A).

We found that the presence of Csy4 generated functional gRNA1, as determined by EYFP activation (Figure 2B-D and Figure S1B for raw data). gRNA1 generated from the HSV1 intron produced the strongest EYFP activation (Figure 2D), which reached saturation at 200 ng of the Csy4 plasmid. In contrast, the snoRNA2 intron saturated EYFP expression at 50 ng of the Csy4 plasmid but the maximal EYFP levels produced by this intron were the lowest of all introns tested (∼65% of the HSV1 intron). In addition, increased Csy4 levels concomitantly reduced mKate2 levels. While these trends were similar for all three introns examined, the magnitudes of the effects were intron-specific. The snoRNA2 intron exhibited the largest decrease in mKate2 levels with increasing Csy4 plasmid concentrations, with a 15-fold reduction in mKate2 fluorescence at 400 ng of the Csy4 plasmid compared to the no Csy4 condition (Figure 2C). The consensus and HSV1 introns exhibited mKate2 levels that were less sensitive to increasing Csy4 levels (Figure 2B and Figure 2D). Thus, together with the ‘triplex/Csy4’ architecture, the ‘intron/Csy4’ approach provides a set of parts for the tunable production of functional gRNAs from translated genes. Specifically, absolute protein levels of the gRNA-harboring genes and downstream target genes, as well as the ratios between them, can be determined by the choice of specific parts and concentration of Csy4.

### Interactions between Csy4 and intronic gRNA

To determine whether both of the 5’ and 3’ Csy4 recognition sites are necessary for functional gRNA generation from introns, we used an HSV1-based intron within mKate2. This intron housed a gRNA1 sequence that was either preceded by a Csy4 binding site on its 5’ side (‘28-gRNA’, Figure 2E and Figure S4) or followed by a Csy4 binding site on its 3’ end (‘gRNA-28’, Figure 2F and Figure S4). The synthetic P1-EYFP construct was used to assess gRNA1 activity. The data for Figure 2E,F was normalized with the performance of the ‘intron/Csy4’ architecture where intronic gRNA1 was flanked by two Csy4 binding sites (’28-gRNA-28’, Figure S4). Both architectures containing only a single Csy4 binding site had mKate2 levels which decreased with the addition of Csy4 versus no Csy4 (Figure 2E,F).

In contrast, downstream EYFP activation by the gRNA1-directed CRISPR-TF was significantly lower for the single Csy4-binding-site architectures (Figure 2E,F) versus the ‘intron/Csy4’ construct (Figure 2D). When only one Csy4 binding site was located at the 5’ end of the gRNA1 intron, EYFP expression was not detectable (Figure 2E). When only one Csy4 binding site was located at the 3’ end of the gRNA1 intron, a 6-fold reduction in EYFP levels was observed (Figure 2F) compared with the ‘intron/Cys4’ architecture which contains Csy4 recognition sites flanking gRNA1 (Figure 2D). These results demonstrate that the 5’ Csy4 recognition sequence is important for generating functional intronic gRNAs and the 3’ Csy4 binding site is essential. We hypothesize that Csy4 can help stabilize intronic gRNA through several potential mechanisms which can be investigated in future work. The 5’ end of RNAs cleaved by Csy4 contain a hydroxyl (OH^-^) which may protect them from major 5’→3’ cellular RNAases such as the XRN family, which require a 5’ phosphate for substrate recognition (Houseley and Tollervey, 2009; Nagarajan et al., 2013). In addition, binding of the Csy4 protein to the 3’ end of the cleaved gRNA (Haurwitz et al., 2012) may protect it from 3’→5’ degradation mediated by the eukaryotic exosome complex (Houseley and Tollervey, 2009).

### Functional gRNA generation with *cis*-acting ribozymes

Very recently, the generation of gRNAs from RNAP II promoters for genome editing applications has been demonstrated in wheat (Upadhyay et al., 2013) and in yeast (Gao and Zhao, 2014) by using *cis*-acting ribozymes. In addition to the ‘triplex/Csy4’ and ‘intron/Csy4’-based mechanisms described above, we also employed self-cleaving ribozymes to enable gene regulation in human cells via gRNAs generated from RNAP II promoters. Specifically, our gRNAs were engineered to contain a hammerhead (HH) ribozyme (Pley et al., 1994) on their 5’ end and a HDV ribozyme (Ferre-D’Amare et al., 1998) on their 3’ end, as shown in Figure 3.

We tested ribozymes in three different architectures, all driven by a CMVp: (1) an mKate2 transcript followed by a triplex and a HH-gRNA1-HDV sequence (CMVp-mK-Tr-HH-g1-HDV, Figure 3A); (2) an mKate2 transcript followed a HH-gRNA1-HDV sequence (CMVp-mK-HH-g1-HDV, Figure 3B); and (3) the sequence HH-gRNA1-HDV itself with no associated protein coding sequence (CMVp-HH-g1-HDV, Figure 3C). We compared gRNAs generated from these architectures with gRNAs produced by the RNAP III promoter U6 and the ‘triplex/Cys4’ architecture (with 200 ng of the Csy4 plasmid) described earlier. All constructs utilized gRNA1, which drove the expression of EYFP from a P1-EYFP-containing plasmid.

**Figure 3.**
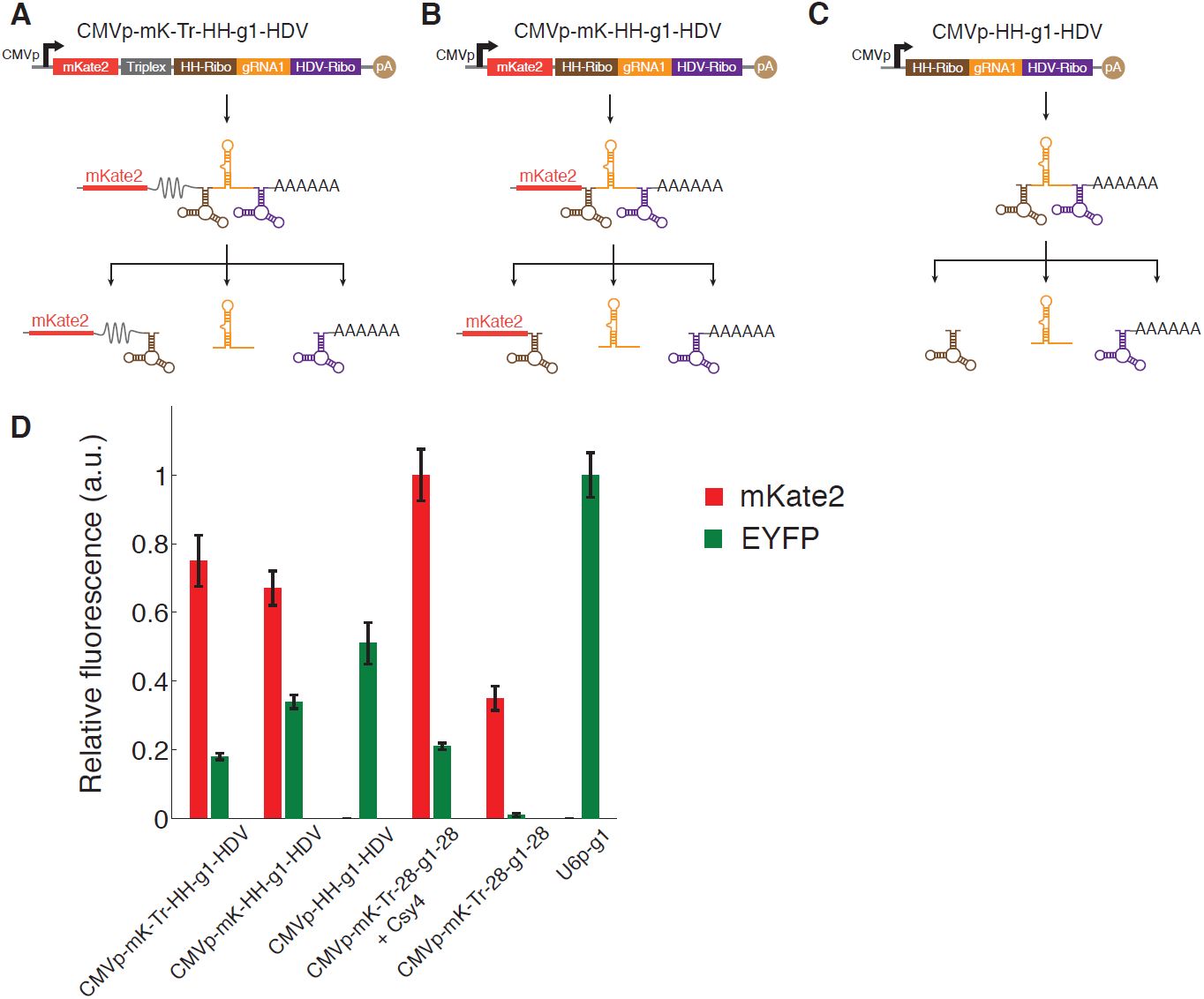
Ribozyme architectures can produce active gRNAs while maintaining expression of the harboring gene. **(A)** gRNA1 was flanked with a hammerhead (HH) ribozyme and an HDV ribozyme. This construct was encoded downstream of mKate2 with an RNA triplex and expressed from the CMV promoter (CMVp-mK-Tr-HH-g1-HDV). **(B)** gRNA1 was flanked with a hammerhead (HH) ribozyme and an HDV ribozyme. This construct was encoded downstream of mKate2 with no RNA triplex and expressed from the CMV promoter (mK-HH-g1-HDV). **(C)** gRNA1 was flanked with a hammerhead (HH) ribozyme and an HDV ribozyme. This construct was expressed from the CMV promoter (HH-g1-HDV). **(D)** Functional gRNA1 generation was assessed by EYFP expression resulting from the activation of P1-EYFP. The three ribozyme-based architectures were able to efficiently activate EYFP expression. The ‘triplex/Csy4’ construct (mK-Tr-28-g1-28), with and without Csy4, as well as the RNAP III promoter U6p driving gRNA1 (U6p-g1) are shown for comparison.

All the constructs that contained mKate2 exhibited detectable mKate2 fluorescence levels (Figure 3D and Figure S5). Surprisingly, this included CMVp-mK-HH-g1-HDV, which did not have a triplex sequence and was thus expected to have low mKate2 levels due to removal of the poly-(A) tail. This could be due to inefficient ribozyme cleavage (Beck and Nassal, 1995; Chowrira et al., 1994; R Hormes, 1997) which allows non-processed transcripts to be transported to the cytoplasm and translated, protection of the mKate2 transcript by the residual 3’ ribozyme sequence, or other mechanisms, but is yet to be determined. In terms of output EYFP activation, the highest EYFP fluorescence level was generated from gRNAs expressed by U6p, followed by the CMVp-HH-g1-HDV and CMVp-mK-HH-g1-HDV constructs (Figure 3D). The CMVp-mK-Tr-HH-gRNA1-HDV and ‘triplex/Csy4’ architectures had similar EYFP levels.

C*is-*acting ribozymes are useful and can mediate functional gRNA expression from RNAP II promoters. However, these *cis*-ribozyme-based strategies are not readily amenable to synchronization, induction, or rewiring of RNA-directed circuits at the single-cell level. Ribozymes whose activities can be regulated with external ligands, such as theophylline, have been previously described (Koizumi et al., 1999; Soukup and Breaker, 1999) and could be used to trigger gRNA release exogenously. However, such strategies cannot link intracellular ribozyme activity to endogenous signals generated within single cells. In contrast, as shown below, the expression of genetically encoded Csy4 can be used to rewire RNA-directed genetic circuits and change their behavior (Figure 7). In future work, *trans*-activating ribozymes could potentially be used to link RNA cleavage and gRNA generation to intracellular events (Kuwabara et al., 1998).

### Multiplexed gRNA expression from single RNA transcripts

A major challenge in constructing CRISPR-TF-based circuits in human cells, especially ones which interface with endogenous promoters, is that multiple gRNAs are often necessary to achieve desired activation levels (Cheng et al., 2013; Maeder et al., 2013a; Mali et al., 2013a; Perez-Pinera et al., 2013a). Current techniques rely on the use of multiple gRNA expression cassettes, each with their own promoter. The toolkit described here can be used to express many functional gRNAs from a single transcript, thus enabling compact encoding of synthetic gene circuits with multiple outputs as well as concise strategies for modulating native genes.

To this end, we sought to demonstrate the expression of two independent gRNAs from a single RNA transcript to activate two independent downstream promoters. In the first architecture (‘intron-triplex’), we encoded gRNA1 within an HSV1 intron flanked by two Csy4 binding sites within the coding sequence of mKate2. Furthermore, we encoded gRNA2 enclosed by two Csy4 binding sites downstream of the mKate2-triplex sequence (Figure 4A, CMVp-mK_EX1_-[28-g1-28]_HSV1_-mK_EX2_-Tr-28-g2-28). In the second architecture (‘triplex-tandem’), we surrounded both gRNA1 and gRNA2 with Csy4 binding sites and placed them in tandem, downstream of the mKate2-triplex sequence (Figure 4B, CMVp-mK-Tr-28-g1-28-g2-28). In both architectures, gRNA1 and gRNA2 targeted the synthetic promoters P1-EYFP and P2-ECFP, respectively.

**Figure 4.**
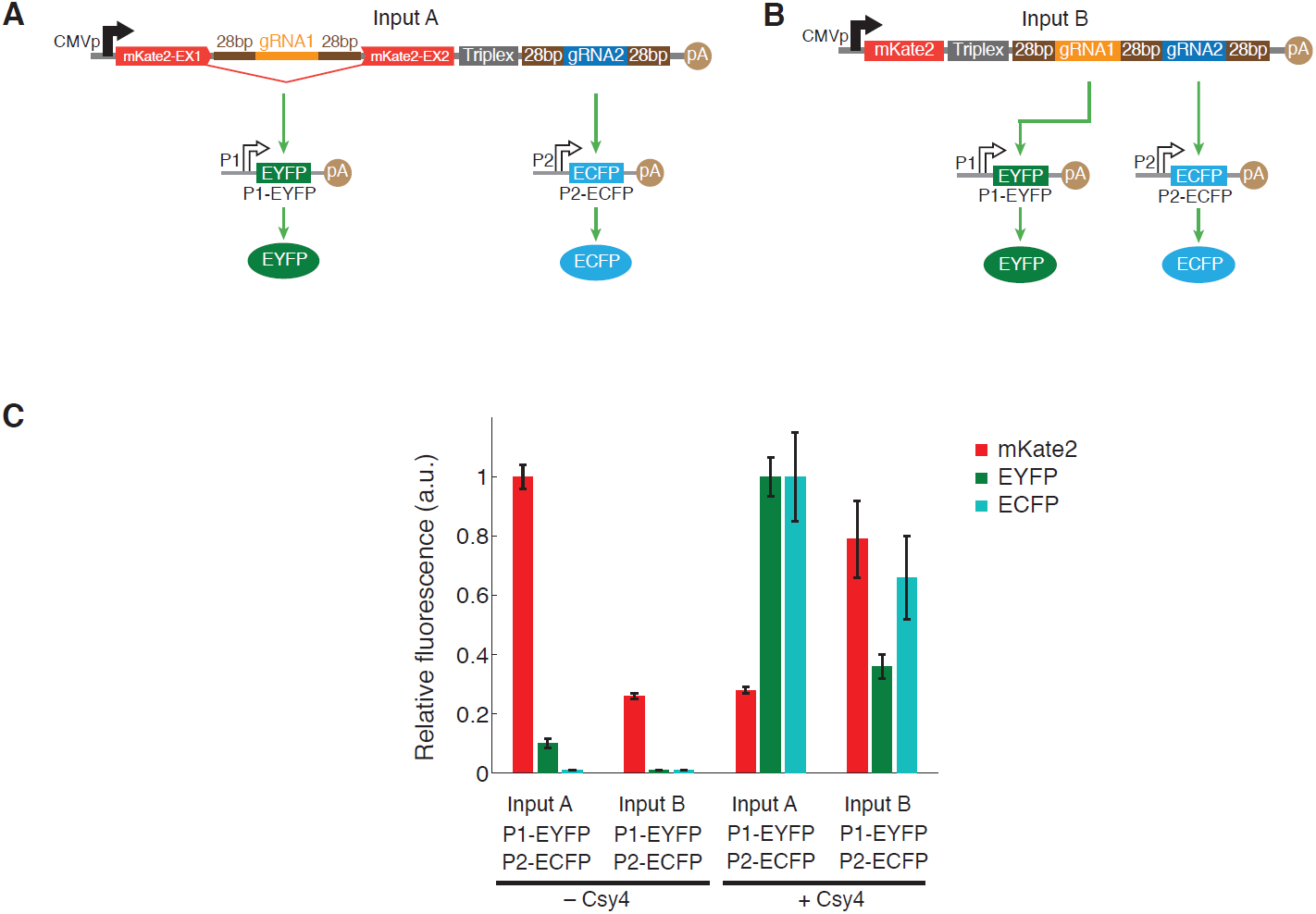
Multiplexed gRNA expression from a single transcript can be achieved with the ‘triplex/Csy4’ and ‘intron/Csy4’ architectures, thus enabling compact encoding of synthetic circuits with multiple outputs. **(A)** In the first design (Input A, ‘intron-triplex’), we encoded gRNA1 within a HSV1 intron and gRNA2 after an RNA triplex. Both gRNAs were flanked by Csy4 recognition sites. Functional gRNA expression was assessed by activation of a gRNA1-specific P1-EYFP construct and a gRNA2-specific P2-ECFP construct. **(B)** In the second design (Input B, ‘triplex tandem’), we encoded both gRNA1 and gRNA2 in tandem, with intervening and flanking Csy4 recognition sites, downstream of mKate2 and an RNA triplex. Functional gRNA expression was assessed by activation of a gRNA1-specific P1-EYFP construct and a gRNA2-specific P2-ECFP construct. **(C)** Both multiplexed gRNA expression constructs exhibited efficient activation of EYFP and ECFP expression in the presence of Csy4, thus demonstrating the generation of multiple active gRNAs from a single transcript. Furthermore, as expected from Figure 1 and Figure 2, mKate2 levels decreased in the first design due to the intronic architecture whereas mKate2 levels increased in the second design due to the non-intronic architecture.

As shown in Figure 4C (see Figure S6 for raw data), both strategies resulted in active multiplexed gRNA production. The ‘intron-triplex’ construct exhibited a 3-fold decrease in mKate2, a 10-fold increase in EYFP, and a 100-fold increase in ECFP in the presence of 200 ng of the Csy4 plasmid compared to no Csy4. In the ‘triplex-tandem’ architecture, mKate2, EYFP, and ECFP expression increased by 3-fold, 36-fold, and 66-fold, respectively, in the presence of 200 ng of the Csy4 plasmid compared to no Csy4. The ‘intron-triplex’ architecture had higher EYFP and ECFP levels compared with ‘triplex-tandem’ construct. Thus, both strategies for multiplexed gRNA expression enable functional CRISPR-TF activity at multiple downstream targets and can be tuned for desired applications.

To further explore the scalability of our multiplexing toolkit and to demonstrate its utility in targeting endogenous loci, we generated four different gRNAs species from a single transcript. The four gRNAs required for IL1RN activation were cloned in tandem, separated by Csy4 binding sites, downstream of an mKate2-triplex sequence on a single transcript (Figure 5A). We compared IL1RN activation by the multiplexed single-transcript construct with an architecture where the four different gRNAs were expressed from four different plasmids (Figure 5B, ‘Multiplexed’ versus ‘Non multiplexed’, respectively). In the presence of 100 ng of the Csy4 plasmid, the multiplexed architecture resulted in a ∼1111-fold activation over non-specific gRNA1 (‘NS’) and was ∼2.5 times more efficient than the non-multiplexed set of single-gRNA-expressing plasmids. Furthermore, ∼155-fold IL1RN activation was detected with the multiplexed architecture even in the absence of Csy4, which suggests that taCas9 can bind to gRNAs and recruit them for gene activation despite no Csy4 being present. These results demonstrate that it is possible to encode multiple functional gRNAs for multiplexed expression from a single concise RNA transcript. These architectures therefore enable compact programming of Cas9 function for implementing multi-output synthetic gene circuits, for modulating endogenous genes, and for potentially achieving conditional multiplexed genome editing.

**Figure 5.**
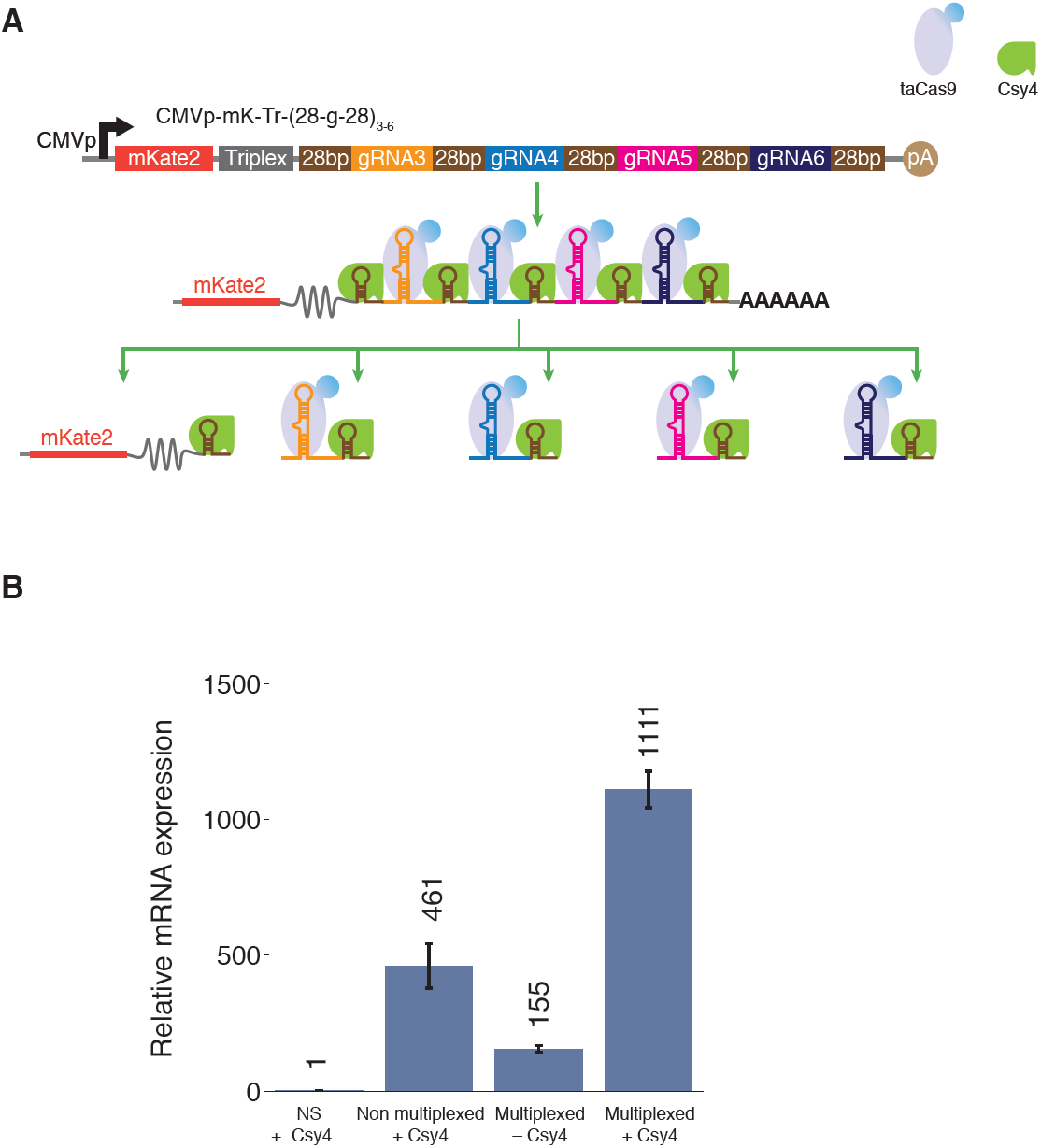
Multiplexed gRNA expression from a single transcript enables efficient activation of endogenous loci with a compact architecture. **(A)** Four different gRNAs (gRNA3-6) were encoded in tandem, with intervening and flanking Csy4 recognition sites, down-stream of mKate2 and an RNA triplex (mK-Tr-(28-g-28)_3-6_). **(B)** The multiplexed mK-Tr-(28-g-28)_3-6_ construct exhibited high-level activation of IL1RN expression in the presence of Csy4 compared with the same construct in the absence of Csy4. Relative IL1RN mRNA expression was determined compared to a control construct with non-specific gRNA1 (NS, CMVp-mK-Tr-28-g1-28) expressed via the ‘triplex/Csy4’ architecture. For comparison, a non-multiplexed set of plasmids containing the same gRNAs (gRNA3-6), each expressed from separate, individual plasmids is shown.

### Synthetic transcriptional cascades with RNA-guided regulation

To demonstrate the utility of our RNA-dependent regulatory toolkit, we used it to create the first CRISPR-TF-based transcriptional cascades. Many complex gene circuits require the ability to implement cascades, in which signals integrated at one stage are transmitted into multiple downstream stages for processing and actuation (Ellis et al., 2009; Hooshangi et al., 2005). For example, gene cascades are important for synthetic-biology applications such as multi-layer artificial gene circuits that compute in living cells (Weber and Fussenegger, 2009). Furthermore, transcriptional cascades are important in natural regulatory systems, such as those that control segmentation, sexual commitment, and development (Dequeant and Pourquie, 2008; Peel et al., 2005; Sinha et al., 2014). However, despite the potential utility of CRISPR-TFs for artificial gene circuits, CRISPR-TF-based cascades have not been built to-date.

We integrated the ‘triplex/Csy4’ and ‘intron/Csy4’ strategies to build two different three-stage CRISPR-TF-mediated transcriptional cascades (Figure 6). In the first design, CMVp-driven expression of gRNA1 from an ‘intron/Csy4’ construct generated gRNA1 from an HSV1 intron, which activated a synthetic promoter P1 to produce gRNA2 from a ‘triplex/Csy4’ architecture, which then activated a downstream synthetic promoter P2 regulating ECFP (Figure 6A). In the second design, the intronic gRNA expression cassette in the first stage of the cascade was replaced by a ‘triplex/Csy4’ architecture for expressing gRNA1 (Figure 6B). We tested these two designs in the presence of 200 ng of the Csy4 plasmid (Figure 6C,D and Figure S7).

**Figure 6.**
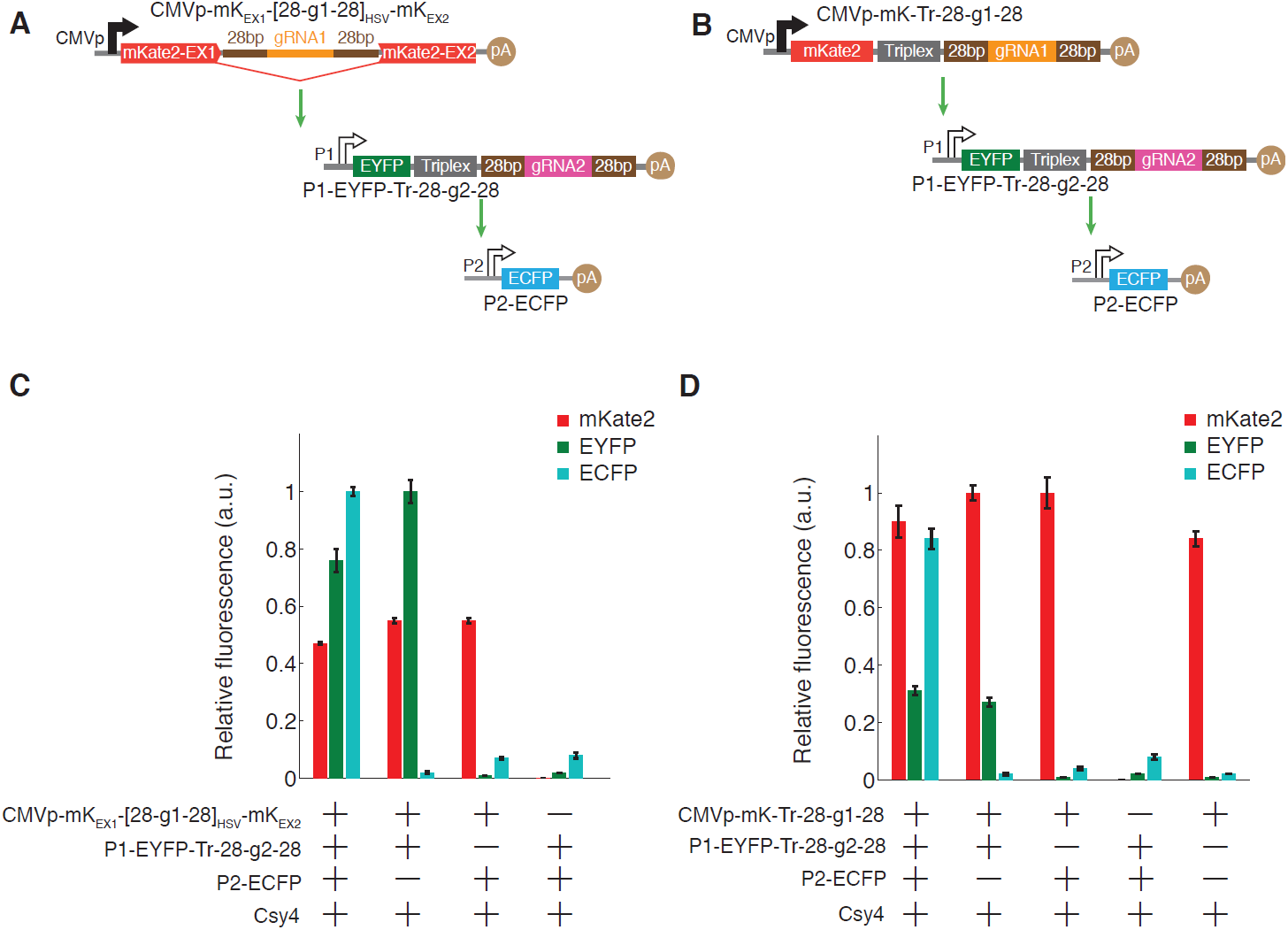
Multi-layer regulation is important for sophisticated behaviors in artificial and natural gene circuits. Multi-stage transcriptional cascades can be implemented with CRISPR-TFs using the architectures described here. **(A)** A three-stage transcriptional cascade was implemented by using intronic gRNA1 (CMVp-mK_EX1_-[28-g1-28]_HSV_-mK_EX2_) as the first stage. gRNA1 specifically targeted the P1 promoter to express gRNA2 (P1-EYFP-Tr-28-g2-28), which then activated expression of ECFP from the P2 promoter (P2-ECFP). **(B)** A three-stage transcriptional cascade was implemented by using a ‘triplex/Csy4’ architecture to express gRNA1 (CMVp-mK-Tr-28-g1-28). gRNA1 specifically targeted the P1 promoter to express gRNA2 (P1-EYFP-Tr-28-g2-28), which then activated expression of ECFP from P2 (P2-ECFP). **(C)** The complete three-stage transcrip-tional cascade from A) exhibited expression of all three fluorescent proteins. The removal of one of each of the three stages in the cascade resulted in the expected loss of fluorescence of the specific stage and dependent downstream stages. **(D)** The complete three-stage transcriptional cascade from B) exhibited expression of all three fluorescent proteins. The removal of one of each of the three stages in the cascade resulted in the expected loss of fluorescence of the specific stage and dependent downstream stages.

In the first cascade design, a 76-fold increase in EYFP and a 13-fold increase in ECFP were observed compared to a control in which the second stage of the cascade (P1-EYFP-Tr-28-g2-28) was replaced by an empty plasmid (Figure 6C). In the second cascade design, a 31-fold increase in EYFP and a 21-fold increase in ECFP were observed compared to a control in which the second stage of the cascade (P1-EYFP-Tr-28-g2-28) was replaced by an empty plasmid (Figure 6D). These results demonstrate that there is minimal non-specific activation of promoter P2 by gRNA1, which is essential for the scalability and reliability of transcriptional cascades. Furthermore, the fold-activation of each stage in the cascade was dependent on the presence of all upstream nodes, which is expected in properly functioning transcriptional cascades (Figure 6C,D).

### Rewiring RNA-dependent synthetic regulatory circuits

We next sought to demonstrate how CRISPR-TF regulation can be integrated with mammalian RNA interference to implement more sophisticated circuit topologies. Furthermore, we showed how network motifs could be rewired based on Csy4-based RNA processing. Specifically, we incorporated miRNA regulation with CRISPR-TFs and used Csy4 to disrupt miRNA inhibition of target RNAs by removing cognate miRNA binding sites. We first built a single RNA transcript that was capable of expressing both a functional miRNA (Greber et al., 2008; Xie et al., 2011) and a functional gRNA. This was achieved by encoding a mammalian miRNA inside the consensus intron within the mKate2 gene, followed by a triplex sequence and a gRNA1 sequence flanked by Csy4 recognition sites (Figure 7A, CMVp-mK_Ex1_-[miR]-mK_Ex2_-Tr-28-g1-28). We also implemented two output constructs to demonstrate the potential for multiplexed gene regulation with our toolkit. The first output was a constitutively expressed ECFP gene followed by a triplex sequence, a Csy4 recognition site, 8x miRNA binding sites (8x miRNA-BS), and another Csy4 recognition site (Figure 7A). The second output was a synthetic P1 promoter regulating EYFP expression (Figure 7A).

**Figure 7.**
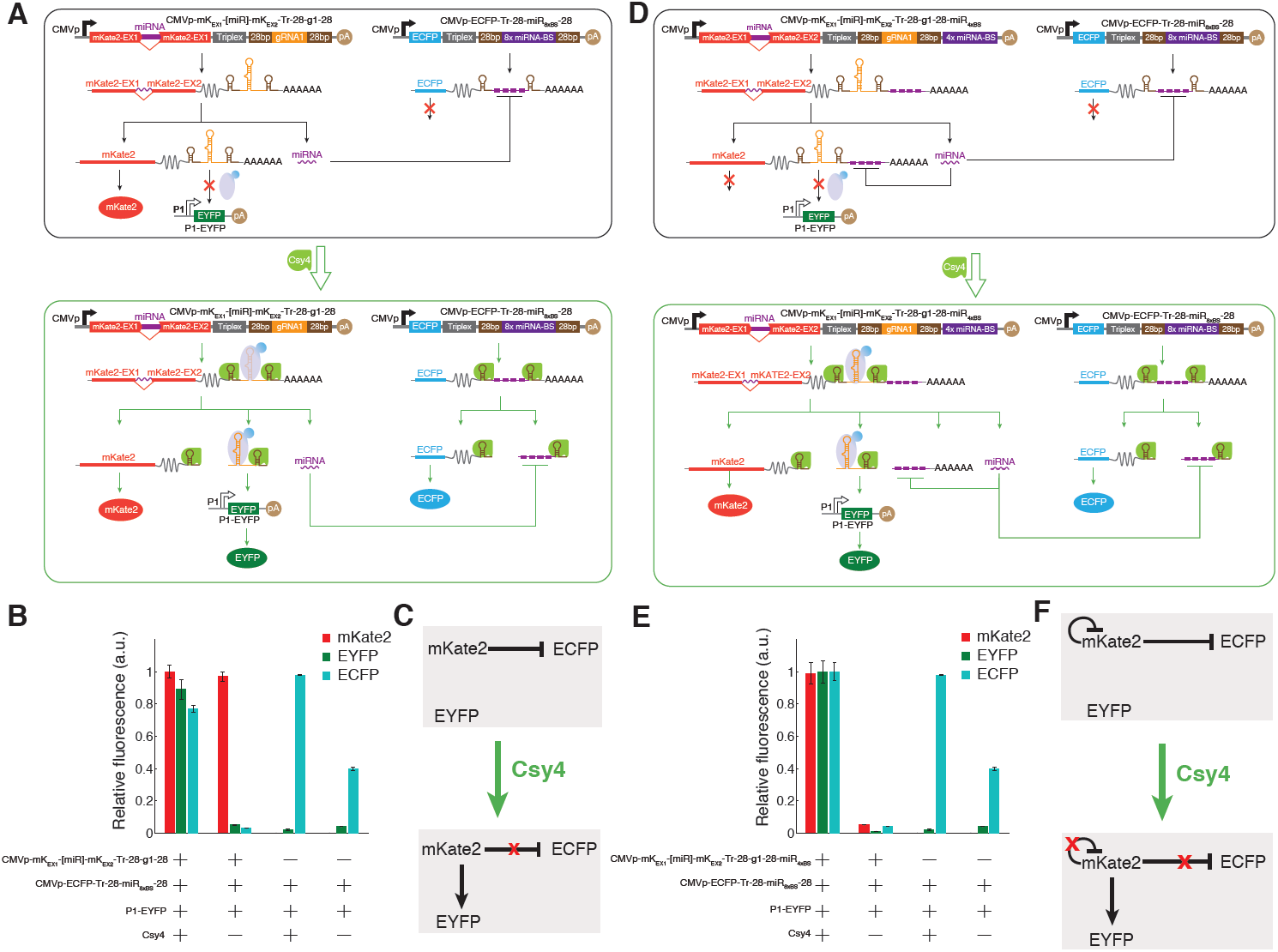
CRISPR-based transcriptional regulation can be integrated with mammalian microRNAs and RNA processing mechanisms as well as with Csy4-dependent RNA processing to implement feedback loops and multi-output circuits that can be rewired at the RNA level. **(A)** We created a single transcript that encoded both miRNA and CRISPR-TF-based regulation by expressing a miRNA from an intron within mKate2 and gRNA1 from a ‘triplex/Csy4’ architecture (CMVp-mK_Ex1_-[miR]-mK_Ex2_-Tr-28-g1-28). In the presence of taCas9 but in the absence of Csy4, this circuit did not activate a downstream gRNA1-specific P1-EYFP construct and did repress a downstream ECFP transcript with 8x miRNA binding sites flanked by Csy4 recognition sites (CMVp-ECFP-Tr-28-miR_8xBS_). In the presence of both taCas9 and Csy4, this circuit was rewired by activating gRNA1 production and subsequent EYFP expression as well as by separating the ECFP transcript from the 8xmiRNA binding sites, thus ablating miRNA inhibition of ECFP expression. **(B-C)** The results demonstrate that Csy4 expression can change the behavior of the circuit in A) by rewiring circuit interconnections. A circuit motif diagram illustrates the Csy4-catalyzed rewiring. **(D)** We incorporated an autoregulatory feedback loop into the network topology of the circuit described in A) by encoding 4x miRNA binding sites at the 3’ end of the input transcript (CMVp-mK_Ex1_-[miR]-mK_Ex2_-Tr-28-g1-28-miR_4xBS_). This negative feedback suppressed mKate2 expression in the absence of Csy4. However, in the presence of Csy4, the 4x miRNA binding sites were separated from the mKate2 mRNA, thus leading to mKate2 expression. **(E-F)** The results demonstrate that Csy4 expression can change the behavior of the circuit in D) by rewiring circuit interconnections. In contrast to the circuit in A), mKate2 was suppressed in the absence of Csy4 but was highly expressed in the presence of Csy4 due to elimination of the miRNA-based autoregulatory negative feedback. **(F)** A circuit motif diagram illustrates the Csy4-catalyzed rewiring. Each of the mKate, EYFP, and ECFP levels in B) and E) were normalized to the respective maximal fluorescence levels amongst all the tested scenarios. We note that the controls in column 3 and 4 in B) and E) are duplicated, as the two circuits in A) and D) were tested in the same experiment with the same controls.

In the absence of Csy4, ECFP and EYFP levels were low because the miRNA suppressed ECFP expression and no functional gRNA1 was generated (Figure 7B and Figure S8 ‘Mechanism 1’). In the presence of Csy4, ECFP expression increased by 30-fold compared to the no Csy4 condition, which we attributed to Csy4-induced separation of the 8x miRNA-BS from the ECFP transcript (Figure 7B). Furthermore, the presence of Csy4 generated functional gRNA1, leading to 17-fold increased EYFP expression compared to the no Csy4 condition (Figure 7B). The mKate2 fluorescence levels were high in both the Csy4-positive and Csy4-negative conditions. Thus, Csy4 catalyzed RNA-based rewiring of circuit connections between the input node and its two outputs by simultaneously inactivating a repressive output link and enabling an activating output link (Figure 7C).

To demonstrate the facile nature by which additional circuit topologies can be programmed using RNA-dependent mechanisms, we extended the design in Figure 7A by incorporating an additional 4x miRNA-BS at the 3’ end of the mKate-containing transcript (Figure 7D, CMVp-mK_Ex1_-[miR]-mK_Ex2_-Tr-28-g1-28-miR_4xBS_). In the absence of Csy4, this resulted in autoregulatory negative-feedback suppression of mKate2 expression by the miRNA generated within the mKate2 intron (Figure 7E and Figure S8 ‘Mechanism 2’). In addition, both ECFP and EYFP levels remained low due to repression of ECFP by the miRNA and the lack of functional gRNA1 generation. However, in the presence of Csy4, mKate2 levels increased by 21-fold due to Csy4-mediated separation of the 4x miRNA-BS from the mKate2 transcript. Furthermore, ECFP inhibition by the miRNA was relieved in a similar fashion, resulting in a 27-fold increase in ECFP levels. Finally, functional gRNA1 was generated, leading to a 50-fold increase in EYFP levels (Figure 7E). Thus, Csy4 catalyzed RNA-based rewiring of circuit connections between the input node and its two outputs by simultaneously inactivating a repressive output link, enabling an activating output link, and inactivating an autoregulatory feedback loop (Figure 7F).

In summary, we have shown that our RNA-dependent regulatory toolkit facilitates the construction of multi-mechanism genetic circuits that integrate miRNAs and CRISPR-TFs for tunable, multi-output gene regulation. Furthermore, Csy4-based RNA processing can be used to rewire multiple interconnections and feedback loops between genetic components, resulting in synchronized shifts in circuit behavior.

## Summary

Synthetic biology provides tools for studying natural regulatory networks by disrupting, rewiring, and mimicking natural network motifs (Bashor et al., 2008; Deans et al., 2007; Elowitz and Lim, 2010; Guido et al., 2006; Kampf et al., 2012; Kemmer et al., 2011; Levine et al., 2012; McMillen et al., 2002; Nissim et al., 2007; Park et al., 2003; Sprinzak et al., 2011). In addition, synthetic circuits have been used to link exogenous signals to endogenous gene regulation (Fussenegger et al., 2000; Muller et al., 2013a; Muller et al., 2013b; Weber et al., 2007; Weber et al., 2002; Ye et al., 2011), to address biomedical applications (Kemmer et al., 2010; Nissim and Bar-Ziv, 2010; Weber and Fussenegger, 2012; Weber et al., 2008; Ye et al., 2011), and to perform cellular computation (Benenson, 2012; Rinaudo et al., 2007; Tabor et al., 2009). Although many synthetic gene circuits have been based on transcriptional regulation, RNA-based regulation has also recently been used to construct a variety of synthetic gene circuits (Auslander et al., 2012; Babiskin and Smolke, 2011; Chen et al., 2010; Culler et al., 2010; Delebecque et al., 2011; Kennedy et al., 2013; Rinaudo et al., 2007; Saito et al., 2011; Saito et al., 2010; Win et al., 2009; Xie et al., 2011). However, despite many advances, previous efforts have not yet integrated RNA-based regulation with CRISPR-TFs, which are both promising strategies for implementing scalable genetic circuits given their programmability and potential for multiplexing. In this work, we created a rich toolkit for engineering artificial gene circuits and endogenous gene regulation in human cells. This framework integrates mammalian RNA regulatory mechanisms with the RNA-dependent protein, dCas9, and the RNA-processing protein, Csy4, from bacteria. Moreover, it enables convenient programming of regulatory links based on base-pairing complementary between nucleic acids.

We developed multiple complementary approaches to generate functional gRNAs from the coding sequence of proteins regulated by RNAP II promoters, which also permit concomitant expression of the harboring gene. The harboring genes we used were fluorescent genes, since they are convenient reporters of promoter activity. However, these genes can be readily swapped with any other protein-coding sequence, thus enabling multiplexed expression of gRNAs along with arbitrary protein outputs from a single construct. We validated the ability of these strategies, based on RNA triplexes with Csy4, RNA introns with Csy4, and *cis*-acting ribozymes, to generate functional gRNAs by targeting synthetic promoters. Furthermore, when gRNAs were flanked by Csy4 recognition sites and located downstream of a harboring gene followed by an RNA triplex, the levels of the harboring gene increased with the levels of Csy4. The opposite effect was found when gRNAs were flanked by Csy4 recognition sites within introns, with the magnitude of the effect varying depending on the specific intronic sequence used. Thus, these complementary architectures enable tunable RNA and protein levels to be achieved within synthetic gene circuits.

As a complement to synthetic circuits, we showed that this toolkit can be used to activate endogenous promoters from multiple different human RNAP II promoters, as well as the CMV promoter. Importantly, we described novel strategies for multiplexed gRNA expression from compact single transcripts to modulate both synthetic and native promoters. This feature is useful because it can be used to regulate multiple nodes from a single one. The ability to concisely encode multiple gRNAs within a single transcript enables sophisticated circuits with a large number of parallel ‘fan-outs’ (i.e., outgoing interconnections from a given node) and networks with dense interconnections. Moreover, the ability to synergistically modulate endogenous loci with several gRNAs in a condensed fashion is critical since multiple gRNAs are often needed to enact substantial modulation of native promoters (Perez-Pinera et al., 2013a). Thus, the tools described here can be used to build efficient artificial gene networks and to perturb native regulatory networks.

It has been previously shown that the native CRISPR RNA context can be used to perform multiplexed genome engineering when expressed from RNAP III promoters in mammalian cells (Cong et al., 2013). However, it remains unclear whether CRISPR/Cas-based systems can be multiplexed when expressed from RNAP II promoters using this approach and what cellular factors mediate this process. It has also been shown that individual gRNA-expressing plasmids can be co-transfected to enable multiplexed genome editing (Yang et al., 2013). In addition to transcriptional regulation, if a nuclease-proficient Cas9 was used with our platform instead of taCas9, then multiplexed genome-editing activity could be conditionally linked to cellular signals via regulation of gRNA expression. This would enable conditional, multiplexed knockouts within *in vivo* settings – for example, with cell-specific, temporal, or spatial control. In addition to genetic studies, this capability could be potentially used to create *in vivo* DNA-based ‘ticker tapes’ that link cellular events to mutations.

These architectures lay down a foundation for the construction of sophisticated and compact synthetic gene circuits in human cells. Theoretically, since the specificity of regulatory interconnections with these tools is determined only by RNA sequences, scalable circuits with almost any network topology could be constructed. For example, multi-layer network topologies are important for achieving sophisticated behaviors, both in artificial and natural genetic contexts (Alon, 2007). Thus, to demonstrate the utility of our toolkit for implementing more complex synthetic circuits, we used it to create the first CRISPR-TF-based transcriptional cascades which were highly specific and effective. We demonstrated reliable three-step transcriptional cascades with two different architectures that incorporated RNA triplexes, introns, Csy4, and CRISPR-TFs. The absence of undesired crosstalk between different stages of the cascade underscores the orthogonality and scalability of RNA-dependent regulatory schemes for synthetic gene circuit design. Combining multiplexed gRNA expression with transcriptional cascades could be used to create multi-stage, multi-input/multi-output gene networks capable of logic, computing, and interfacing with endogenous systems. In addition, useful topologies, such as multi-stage feedforward and feedback loops, can be readily programmed.

Furthermore, we showed that RNA regulatory parts, CRISPR-TFs, and RNA interference could be integrated together to create various circuit topologies that can be rewired via conditional RNA processing. Since both positive and negative regulation is possible with the same taCas9 protein (Farzadfard et al., 2013) and miRNAs enact tunable negative regulation, many important multi-component network topologies can be implemented using this set of regulatory parts (Shoval and Alon, 2010). In addition, Csy4 can be used to catalyze changes in gene expression by modifying RNA transcripts. For example, functional gRNAs were liberated for transcriptional modulation and miRNA binding sites were removed from RNA transcripts to eliminate miRNA-based links. In addition, we used the absence or presence of Csy4 to switch a miRNA-based autoregulatory negative feedback loop on and off, respectively (Figure 7B). This feature could be extended in future circuits to minimize unwanted leakage in positive-feedback loops and to dynamically switch circuits between different states. By linking Csy4 expression to endogenous promoters, interconnections between circuits and network behavior could also be conditionally linked to specific tissues, events (e.g., cell cycle phase, mutations), or environmental conditions. With genome mining or directed mutagenesis (Ellefson et al., 2014; Esvelt et al., 2011) on Csy4, orthogonal Csy4 variants could be potentially discovered and used for more complicated RNA processing schemes (Barrangou and van der Oost, 2013). Moreover, additional flexibility and scalability can be achieved by using orthogonal Cas9 proteins (Esvelt et al., 2013).

In summary, this work provides a diverse set of tools for constructing scalable regulatory gene circuits, tuning them, modifying connections between circuit components, and synchronizing the expression of multiple genes in a network. Furthermore, these regulatory parts can be used to interface synthetic gene circuits with endogenous systems as well as to rewire endogenous networks. We envision that integrating RNA-dependent regulatory mechanisms with RNA processing will enable sophisticated transcriptional and post-transcriptional regulation, accelerate synthetic biology, and facilitate the study of basic biology in human cells.

## EXPERIMENTAL PROCEDURES

### Plasmid construction

The **CMVp-dCas9-3xNLS-VP64** (taCas9, Construct 1, Table S1) plasmid was built as described previously (Farzadfard et al., 2013). The *csy4* gene from *Pseudomonas aeruginosa* strain UCBPP-PA14 (Qi et al., 2012), was codon optimized for expression in human cells, PCR amplified to contain an N-terminal 6x-His tag and a TEV recognition sequence, and cloned downstream of the PGK1 promoter between HindIII/SacI sites in the PGK1-EBFP2 plasmid (Farzadfard et al., 2013) to create **PGK1p-Csy4-pA** (Construct 2, Table S1).

The plasmid **CMVp-mKate2-Triplex-28-gRNA1-28-pA** (Construct 3, Table S1) was built using Gibson Assembly from three parts amplified with appropriate homology overhangs: 1) the full length coding sequence of mKate2; 2) the first 110 bp of the mouse MALAT1 3’ triple helix (Wilusz et al., 2012); and 3) gRNA1 containing a 20 bp Specificity Determining Sequence (SDS) and a *S. pyogenes* gRNA scaffold along with 28nt Csy4 recognition sites.

The reporter plasmids **P1-EFYP-pA** (Construct 5, Table S1) and **P2-ECFP-pA** (Construct 6, Table S1) were built by cloning in eight repeats of gRNA1 binding sites and eight repeats of gRNA2 binding sites into the NheI site of **pG5-Luc** (Promega) via annealing complementary oligonucleotides. Then, EYFP and ECFP were cloned into the NcoI/FseI sites, respectively.

The plasmid **CMVp-mKate2_EX1-**[**28-gRNA1-28**]**_HSV1_-mKate2_EX2-pA** (Construct 4, Table S1) was built by Gibson Assembly of the following parts with appropriate homology overhangs: 1) the mKate2_EX1 (a.a 1-90) of mKate2; 2) mKate_EX2 (a.a 91-239) of mKate2; and 3) gRNA1 containing a 20 bp SDS followed by the *S. pyogenes* gRNA scaffold flanked by Csy4 recognition sites and the HSV1 acceptor, donor and branching sequences. Variations of the **CMVp-mKate2_EX1-**[**28-gRNA1-28**]**_HSV1_-mKate2_EX2-pA** plasmid containing consensus and SnoRNA2 acceptor, donor, and branching sequences and with and without the Csy4 recognition sequences (Constructs 8-12, Table S1) were built in a similar fashion.

The ribozyme-expressing plasmids **CMVp-mKate2-Triplex-HHRibo-gRNA1-HDVRibo-pA** and **CMVp-mKate2-HHRibo-gRNA1-HDVRibo-pA** plasmids (Constructs 13 and 14, respectively, Table S1) were built by Gibson Assembly of XmaI-digested **CMVp-mKate2**, and PCR-extended amplicons of gRNA1 (with and without the triplex and containing HHRibo (Gao and Zhao, 2014) on the 5’ end and HDVRibo (Gao and Zhao, 2014) on the 3’ end). The plasmid **CMVp-HHRibo-gRNA1-HDVRibo-pA** (Construct 15, Table S1) was built similarly by Gibson Assembly of SacI-digested **CMVp-mKate2** and a PCR-extended amplicon of gRNA1 containing HHRibo on the 5’ end and HDVRibo on the 3’ end.

The plasmid **CMVp-mKate2_EX1-**[**28-gRNA1-28**]**_HSV1_-mKate2_EX2-Triplex-28-gRNA2-28-pA** (Construct 16, Table S1) was built by Gibson Assembly of the following parts using appropriate homologies: 1) XmaI-digested **CMVp-mKate2_EX1-**[**28-gRNA1-28**]**_HSV1_-mKate2_EX2-pA** (Construct 4, Table S1) and 2) PCR amplified Triplex-28-gRNA2-28 from **CMVp-mKate2-Triplex-28-gRNA1-28-pA** (Construct 3, Table S1).

The plasmid **CMVp-mKate2-Triplex-28-gRNA1-28-gRNA2-28-pA** (Construct 17, Table S1) was built by Gibson Assembly with the following parts using appropriate homologies: 1) XmaI-digested **CMVp-mKate2-Triplex-28-gRNA1-28-pA** (Construct 3, Table S1) and 2) PCR amplified 28-gRNA2-28.

The plasmid **CMVp-mKate2-Triplex-28-gRNA3-28-gRNA4-28-gRNA5-28-gRNA6-28-pA** (Construct 19, Table S1) was constructed using a Golden Gate approach using the Type IIs restriction enzyme, BsaI. Specifically, the IL1RN targeting gRNA3, gRNA4, gRNA5, gRNA6 sequences containing the 20 bp SDSs along with the *S. pyogenes* gRNA scaffold were PCR amplified to contain a BsaI restriction site on their 5’ ends and Csy4 ‘28’ and BsaI restriction sites on their 3’ ends. The PCR amplified products were subjected to 30 alternating cycles of digestion followed by ligation at 37°C and 20°C, respectively. A 540 bp PCR product containing the gRNA3-28-gRNA4-28-gRNA5-28-gRNA6-28 array was amplified and digested with NheI/XmaI and cloned into the **CMVp-mKate2-Triplex-28-gRNA1-28-pA** plasmid (Construct 3, Table S1).

The CMVp-mKate2_EX1-[miRNA]-mKate2_EX2-pA plasmid containing an intronic FF4 (a synthetic miRNA) was received as a gift from Lila Wroblewska. The synthetic FF4 miRNA was cloned into an intron with consensus acceptor, donor and branching sequences between a.a. 90 and 91 of mKate2 to create **CMVp-mKate2_EX1-**[**miRNA**]**-mKate2_EX2-Triplex-28-gRNA1-28-pA** (Construct 20, Table S1) and **CMVp-mKate2_EX1-**[**miRNA**]**-mKate2_EX2-Triplex-28-gRNA1-28-4xFF4BS-pA** (Construct 21, Table S1).

The plasmid **CMVp-ECFP-Triplex-28-8xmiRNA-BS-28-pA** (Construct 22, Table S1) was cloned via Gibson Assembly with the following parts: 1) full length coding sequence of ECFP and 2) 110 nt of the MALAT1 3’ triple helix sequence amplified via PCR extension with oligonucleotides containing eight FF4 miRNA binding sites and Csy4 recognition sequences on both ends.

### Cell culture and transfections

HEK293 T cells were obtained from the American Tissue Collection Center (ATCC) and were maintained in DMEM supplemented with 10% FBS, 1% penicillin-streptomycin, 1% GlutaMAX, non-essential amino acids at 37°C with 5% CO_2_. HEK293 T cells were transfected with FuGENE®HD Transfection Reagent (Promega) according to the manufacturer’s instructions. Each transfection was made using 200,000 cells/well in a 6-well plate. As a control, with 2 µg of a single plasmid in which a CMV promoter regulated mKate2, transfection efficiencies were routinely higher than 90% (determined by flow cytometry performed with the same settings as the experiments). Unless otherwise indicated, each plasmid was transfected at 1 µg/sample. All samples were transfected with taCas9, unless specifically indicated. Cells were processed for flow cytometry or qRT-PCR analysis 72h after transfection.

### Quantitative reverse transcription–PCR (RT-PCR)

The experimental procedure followed was as described in (Perez-Pinera et al., 2013a). Cells were harvested 72h post-transfection. Total RNA was isolated using the RNeasy Plus RNA isolation kit (Qiagen). cDNA synthesis was performed using the qScript cDNA SuperMix (Quanta Biosciences). Real-time PCR using PerfeCTa SYBR Green FastMix (Quanta Biosciences) was performed with the Mastercycler ep realplex real-time PCR system (Eppendorf) with following oligonucleotide primers: IL1RN - forward GGAATCCATGGAGGGAAGAT, reverse TGTTCTCGCTCAGGTCAGTG; GAPDH-forward CAATGACCCCTTCATTGACC, reverse TTGATTTTGGAGGGATCTCG. The primers were designed using Primer3Plus software and purchased from IDT. Primer specificity was confirmed by melting curve analysis. Reaction efficiencies over the appropriate dynamic range were calculated to ensure linearity of the standard curve. We calculated fold-increases in the mRNA expression of the gene of interest normalized to GAPDH expression by the ddCt method. We then normalized the mRNA levels to the non-specific gRNA1 control condition. Reported values are the means of three independent biological replicates with technical duplicates that were averaged for each experiment. Error bars represent standard error of the mean (s.e.m).

### Flow Cytometry

Cells were harvested with trypsin 72h post-transfection, washed with DMEM media and 1xPBS, re-suspended with 1xPBS into flow cytometry tubes and immediately assayed with a Becton Dickinson LSRII Fortessa flow cytometer. At least 50,000 cells were recorded per sample in each data set. The results of each experiment represent data from at least three biological replicates. Error bars are s.e.m. on the weighted median fluorescence values (see Extended Experimental Procedures for detailed information about data analysis).

## ACKNOWLEDGEMENTS

We thank members of the Lu lab for helpful discussions. We thank Ramiz Daniel for valuable inputs. We thank Ky Lowenhaupt for inputs and manuscript editing. In addition, we thank Lila Wroblewska for sharing the mKate2 plasmid expressing intronic miRNA and Jeremy Wilusz and Courtney JnBaptiste for sharing the cGFP_MALAT1_3’ plasmid. Finally, LN would like to thank Adina Binder-Nissim for support. This work was supported by the Defense Advanced Research Projects Agency and the National Institutes of Health (1DP2OD008435 and 1P50GM098792).

## REFERENCES

Alon, U. (2007). Network motifs: theory and experimental approaches. Nat Rev Genet 8, 450–461.

Audibert, A., Weil, D., and Dautry, F. (2002). In vivo kinetics of mRNA splicing and transport in mammalian cells. Mol Cell Biol 22, 6706–6718.

Auslander, S., Auslander, D., Muller, M., Wieland, M., and Fussenegger, M. (2012). Programmable single-cell mammalian biocomputers. Nature 487, 123–127.

Babiskin, A.H., and Smolke, C.D. (2011). A synthetic library of RNA control modules for predictable tuning of gene expression in yeast. Molecular systems biology 7, 471.

Barrangou, R., and van der Oost, J. (2013). RNA-mediated Adaptive Immunity in Bacteria and Archaea (Springer).

Bashor, C.J., Helman, N.C., Yan, S., and Lim, W.A. (2008). Using engineered scaffold interactions to reshape MAP kinase pathway signaling dynamics. Science 319, 1539–1543.

Beck, J., and Nassal, M. (1995). Efficient hammerhead ribozyme-mediated cleavage of the structured heapatitis B virus encapsidation signal in vitro and in cell extracts, but not in intact cells. Nucleic Acids Research 23, 4954–4962.

Beerli, R.R., and Barbas, C.F., 3rd (2002). Engineering polydactyl zinc-finger transcription factors. Nat Biotechnol 20, 135–141.

Benenson, Y. (2012). Biomolecular computing systems: principles, progress and potential. Nat Rev Genet 13, 455–468.

Block, T.M., and Hill, J.M. (1997). The latency associated transcripts (LAT) of herpes simplex virus: still no end in sight. J Neurovirol 3, 313–321.

Blount, B.A., Weenink, T., Vasylechko, S., and Ellis, T. (2012). Rational diversification of a promoter providing fine-tuned expression and orthogonal regulation for synthetic biology. PLoS One 7, e33279.

Chen, M., and Manley, J.L. (2009). Mechanisms of alternative splicing regulation: insights from molecular and genomics approaches. Nature reviews Molecular cell biology 10, 741–754.

Chen, Y.Y., Jensen, M.C., and Smolke, C.D. (2010). Genetic control of mammalian T-cell proliferation with synthetic RNA regulatory systems. Proceedings of the National Academy of Sciences of the United States of America 107, 8531–8536.

Cheng, A.W., Wang, H., Yang, H., Shi, L., Katz, Y., Theunissen, T.W., Rangarajan, S., Shivalila, C.S., Dadon, D.B., and Jaenisch, R. (2013). Multiplexed activation of endogenous genes by CRISPR-on, an RNA-guided transcriptional activator system. Cell Res 23, 1163–1171.

Chowrira, B.M., Pavco, P.A., and McSwiggen, J.A. (1994). In vitro and in vivo comparison of hammerhead, hairpin, and hepatitis delta virus self-processing ribozyme cassettes. Journal of Biological Chemistry 269, 25856–25864.

Clement, J.Q., Qian, L., Kaplinsky, N., and Wilkinson, M.F. (1999). The stability and fate of a spliced intron from vertebrate cells. Rna 5, 206–220.

Cong, L., Ran, F.A., Cox, D., Lin, S., Barretto, R., Habib, N., Hsu, P.D., Wu, X., Jiang, W., Marraffini, L.A., et al. (2013). Multiplex Genome Engineering Using CRISPR/Cas Systems. Science 339, 819–823.

Cronin, C.A., Gluba, W., and Scrable, H. (2001). The lac operator-repressor system is functional in the mouse. Genes & development 15, 1506–1517.

Culler, S.J., Hoff, K.G., and Smolke, C.D. (2010). Reprogramming cellular behavior with RNA controllers responsive to endogenous proteins. Science 330, 1251–1255.

Deans, T.L., Cantor, C.R., and Collins, J.J. (2007). A tunable genetic switch based on RNAi and repressor proteins for regulating gene expression in mammalian cells. Cell 130, 363–372.

Delebecque, C.J., Lindner, A.B., Silver, P.A., and Aldaye, F.A. (2011). Organization of intracellular reactions with rationally designed RNA assemblies. Science 333, 470–474.

Dequeant, M.L., and Pourquie, O. (2008). Segmental patterning of the vertebrate embryonic axis. Nat Rev Genet 9, 370–382.

Ellefson, J.W., Meyer, A.J., Hughes, R.A., Cannon, J.R., Brodbelt, J.S., and Ellington, A.D. (2014). Directed evolution of genetic parts and circuits by compartmentalized partnered replication. Nat Biotech 32, 97–101.

Ellis, T., Wang, X., and Collins, J.J. (2009). Diversity-based, model-guided construction of synthetic gene networks with predicted functions. Nat Biotechnol 27, 465–471.

Elowitz, M., and Lim, W.A. (2010). Build life to understand it. Nature 468, 889–890.

Esvelt, K.M., Carlson, J.C., and Liu, D.R. (2011). A system for the continuous directed evolution of biomolecules. Nature 472, 499–503.

Esvelt, K.M., Mali, P., Braff, J.L., Moosburner, M., Yaung, S.J., and Church, G.M. (2013). Orthogonal Cas9 proteins for RNA-guided gene regulation and editing. Nature Methods 10, 1116–1121.

Farzadfard, F., Perli, S.D., and Lu, T.K. (2013). Tunable and Multifunctional Eukaryotic Transcription Factors Based on CRISPR/Cas. ACS Synthetic Biology 2, 604–613.

Feng, J., Bi, C., Clark, B.S., Mady, R., Shah, P., and Kohtz, J.D. (2006). The Evf-2 noncoding RNA is transcribed from the Dlx-5/6 ultraconserved region and functions as a Dlx-2 transcriptional coactivator. Genes & development 20, 1470–1484.

Ferre-D’Amare, A.R., Zhou, K., and Doudna, J.A. (1998). Crystal structure of a hepatitis delta virus ribozyme. Nature 395, 567–574.

Fussenegger, M., Morris, R.P., Fux, C., Rimann, M., von Stockar, B., Thompson, C.J., and Bailey, J.E. (2000). Streptogramin-based gene regulation systems for mammalian cells. Nature biotechnology 18, 1203–1208.

Gao, Y., and Zhao, Y. (2014). Self-processing of ribozyme-flanked RNAs into guide RNAs in vitro and in vivo for CRISPR-mediated genome editing. Journal of Integrative Plant Biology, n/a-n/a.

Gilbert, Luke A., Larson, Matthew H., Morsut, L., Liu, Z., Brar, Gloria A., Torres, Sandra E., Stern-Ginossar, N., Brandman, O., Whitehead, Evan H., Doudna, Jennifer A., et al. (2013). CRISPR-Mediated Modular RNA-Guided Regulation of Transcription in Eukaryotes. Cell 154, 442–451.

Gossen, M., and Bujard, H. (1992). Tight control of gene expression in mammalian cells by tetracycline-responsive promoters. Proceedings of the National Academy of Sciences 89, 5547–5551.

Greber, D., El-Baba, M.D., and Fussenegger, M. (2008). Intronically encoded siRNAs improve dynamic range of mammalian gene regulation systems and toggle switch. Nucleic Acids Res 36, e101.

Guido, N.J., Wang, X., Adalsteinsson, D., McMillen, D., Hasty, J., Cantor, C.R., Elston, T.C., and Collins, J.J. (2006). A bottom-up approach to gene regulation. Nature 439, 856–860.

Haurwitz, R.E., Sternberg, S.H., and Doudna, J.A. (2012). Csy4 relies on an unusual catalytic dyad to position and cleave CRISPR RNA. Embo j 31, 2824–2832.

Hooshangi, S., Thiberge, S., and Weiss, R. (2005). Ultrasensitivity and noise propagation in a synthetic transcriptional cascade. Proc Natl Acad Sci U S A 102, 3581–3586.

Houseley, J., and Tollervey, D. (2009). The Many Pathways of RNA Degradation. Cell 136, 763–776.

Jackson, R.J. (1993). Cytoplasmic regulation of mRNA function: The importance of the 3′ untranslated region. Cell 74, 9–14.

Jiang, W., Bikard, D., Cox, D., Zhang, F., and Marraffini, L.A. (2013). RNA-guided editing of bacterial genomes using CRISPR-Cas systems. Nat Biotech 31, 233–239.

Jinek, M., Chylinski, K., Fonfara, I., Hauer, M., Doudna, J.A., and Charpentier, E. (2012). A Programmable Dual-RNA–Guided DNA Endonuclease in Adaptive Bacterial Immunity. Science 337, 816–821.

Jinek, M., East, A., Cheng, A., Lin, S., Ma, E., and Doudna, J. (2013). RNA-programmed genome editing in human cells. eLife 2, e00471.

Kampf, M.M., Engesser, R., Busacker, M., Horner, M., Karlsson, M., Zurbriggen, M.D., Fussenegger, M., Timmer, J., and Weber, W. (2012). Rewiring and dosing of systems modules as a design approach for synthetic mammalian signaling networks. Molecular bioSystems 8, 1824–1832.

Kemmer, C., Fluri, D.A., Witschi, U., Passeraub, A., Gutzwiller, A., and Fussenegger, M. (2011). A designer network coordinating bovine artificial insemination by ovulation-triggered release of implanted sperms. Journal of controlled release : official journal of the Controlled Release Society 150, 23–29.

Kemmer, C., Gitzinger, M., Daoud-El Baba, M., Djonov, V., Stelling, J., and Fussenegger, M. (2010). Self-sufficient control of urate homeostasis in mice by a synthetic circuit. Nature biotechnology 28, 355–360.

Kennedy, A.B., Liang, J.C., and Smolke, C.D. (2013). A versatile cis-blocking and trans-activation strategy for ribozyme characterization. Nucleic acids research 41, e41.

Khalil, A., Lu, T.K., Bashor, C., Ramirez, C., Pyenson, N., Joung, J.K., and Collins, J.J. (2012). A Synthetic Biology Framework for Programming Eukaryotic Transcription Functions. Cell 150, 647–658.

Kim, Y.-K., and Kim, V.N. (2007). Processing of intronic microRNAs. The EMBO Journal 26, 775–783.

Koizumi, M., Soukup, G.A., Kerr, J.N., and Breaker, R.R. (1999). Allosteric selection of ribozymes that respond to the second messengers cGMP and cAMP. Nature structural biology 6, 1062–1071.

Kuwabara, T., Warashina, M., Tanabe, T., Tani, K., Asano, S., and Taira, K. (1998). A novel allosterically trans-activated ribozyme, the maxizyme, with exceptional specificity in vitro and in vivo. Mol Cell 2, 617–627.

Lee, J.T. (2012). Epigenetic Regulation by Long Noncoding RNAs. Science 14, 1435–1439.

Levine, J.H., Fontes, M.E., Dworkin, J., and Elowitz, M.B. (2012). Pulsed feedback defers cellular differentiation. PLoS biology 10, e1001252.

Lin, R., Maeda, S., Liu, C., Karin, M., and Edgington, T.S. (2006). A large noncoding RNA is a marker for murine hepatocellular carcinomas and a spectrum of human carcinomas. Oncogene 26, 851–858.

Lohmueller, J.J., Armel, T.Z., and Silver, P.A. (2012). A tunable zinc finger-based framework for Boolean logic computation in mammalian cells. Nucleic Acids Research.

Maeder, M.L., Linder, S.J., Cascio, V.M., Fu, Y., Ho, Q.H., and Joung, J.K. (2013a). CRISPR RNA-guided activation of endogenous human genes. Nat Methods 10, 977–979.

Maeder, M.L., Linder, S.J., Reyon, D., Angstman, J.F., Fu, Y., Sander, J.D., and Joung, J.K. (2013b). Robust, synergistic regulation of human gene expression using TALE activators. Nat Meth 10, 243–245.

Maeder, M.L., Thibodeau-Beganny, S., Sander, J.D., Voytas, D.F., and Joung, J.K. (2009). Oligomerized pool engineering (OPEN): an ‘open-source’ protocol for making customized zinc-finger arrays. Nat Protocols 4, 1471–1501.

Mali, P., Aach, J., Stranges, P.B., Esvelt, K.M., Moosburner, M., Kosuri, S., Yang, L., and Church, G.M. (2013a). CAS9 transcriptional activators for target specificity screening and paired nickases for cooperative genome engineering. Nat Biotechnol 31, 833–838.

Mali, P., Yang, L., Esvelt, K.M., Aach, J., Guell, M., DiCarlo, J.E., Norville, J.E., and Church, G.M. (2013b). RNA-Guided Human Genome Engineering via Cas9. Science 339, 823–826.

McMillen, D., Kopell, N., Hasty, J., and Collins, J.J. (2002). Synchronizing genetic relaxation oscillators by intercell signaling. Proceedings of the National Academy of Sciences 99, 679–684.

Mercer, T.R., Dinger, M.E., and Mattick, J.S. (2009). Long non-coding RNAs: insights into functions. Nat Rev Genet 10, 155–159.

Muller, K., Engesser, R., Metzger, S., Schulz, S., Kampf, M.M., Busacker, M., Steinberg, T., Tomakidi, P., Ehrbar, M., Nagy, F., et al. (2013a). A red/far-red light-responsive bi-stable toggle switch to control gene expression in mammalian cells. Nucleic acids research 41, e77.

Muller, K., Engesser, R., Schulz, S., Steinberg, T., Tomakidi, P., Weber, C.C., Ulm, R., Timmer, J., Zurbriggen, M.D., and Weber, W. (2013b). Multi-chromatic control of mammalian gene expression and signaling. Nucleic Acids Res 41, e124.

Nagano, T., Mitchell, J.A., Sanz, L.A., Pauler, F.M., Ferguson-Smith, A.C., Feil, R., and Fraser, P. (2008). The Air Noncoding RNA Epigenetically Silences Transcription by Targeting G9a to Chromatin. Science 322, 1717–1720.

Nagarajan, V.K., Jones, C.I., Newbury, S.F., and Green, P.J. (2013). XRN 5′->3′ exoribonucleases: Structure, mechanisms and functions. Biochimica et Biophysica Acta (BBA) - Gene Regulatory Mechanisms 1829, 590–603.

Nissim, L., and Bar-Ziv, R.H. (2010). A tunable dual-promoter integrator for targeting of cancer cells. Mol Syst Biol 6, 444.

Nissim, L., Beatus, T., and Bar-Ziv, R. (2007). An autonomous system for identifying and governing a cell’s state in yeast. Physical biology 4, 154–163.

Orioli, A., Pascali, C., Pagano, A., Teichmann, M., and Dieci, G. (2012). RNA polymerase III transcription control elements: Themes and variations. Gene 493, 185–194.

Pandey, R.R., Mondal, T., Mohammad, F., Enroth, S., Redrup, L., Komorowski, J., Nagano, T., Mancini-DiNardo, D., and Kanduri, C. (2008). Kcnq1ot1 Antisense Noncoding RNA Mediates Lineage-Specific Transcriptional Silencing through Chromatin-Level Regulation. Molecular Cell 32, 232–246.

Park, S.H., Zarrinpar, A., and Lim, W.A. (2003). Rewiring MAP kinase pathways using alternative scaffold assembly mechanisms. Science 299, 1061–1064.

Peel, A.D., Chipman, A.D., and Akam, M. (2005). Arthropod segmentation: beyond the Drosophila paradigm. Nat Rev Genet 6, 905–916.

Perez-Pinera, P., Kocak, D.D., Vockley, C.M., Adler, A.F., Kabadi, A.M., Polstein, L.R., Thakore, P.I., Glass, K.A., Ousterout, D.G., Leong, K.W., et al. (2013a). RNA-guided gene activation by CRISPR-Cas9-based transcription factors. Nat Meth 10, 973–976.

Perez-Pinera, P., Ousterout, D.G., Brunger, J.M., Farin, A.M., Glass, K.A., Guilak, F., Crawford, G.E., Hartemink, A.J., and Gersbach, C.A. (2013b). Synergistic and tunable human gene activation by combinations of synthetic transcription factors. Nat Meth 10, 239–242.

Pley, H.W., Flaherty, K.M., and McKay, D.B. (1994). Three-dimensional structure of a hammerhead ribozyme. Nature 372, 68–74.

Proudfoot, N.J. (2011). Ending the message: poly(A) signals then and now. Genes and Development 25, 1770–1782.

Qi, L., Haurwitz, R.E., Shao, W., Doudna, J.A., and Arkin, A.P. (2012). RNA processing enables predictable programming of gene expression. Nat Biotech 30, 1002–1006.

R Hormes, M.H., I Oelze, P Marschall, M Tabler, F Eckstein, and G Sczakiel (1997). The subcellular localization and length of hammerhead ribozymes determine efficacy in human cells. Nucleic Acids Research 25, 769–775.

Reyon, D., Tsai, S.Q., Khayter, C., Foden, J.A., Sander, J.D., and Joung, J.K. (2012). FLASH assembly of TALENs for high-throughput genome editing. Nat Biotech 30, 460–465.

Rinaudo, K., Bleris, L., Maddamsetti, R., Subramanian, S., Weiss, R., and Benenson, Y. (2007). A universal RNAi-based logic evaluator that operates in mammalian cells. Nat Biotechnol 25, 795–801.

Rinn, J.L., Kertesz, M., Wang, J.K., Squazzo, S.L., Xu, X., Brugmann, S.A., Goodnough, L.H., Helms, J.A., Farnham, P.J., Segal, E., et al. (2007). Functional Demarcation of Active and Silent Chromatin Domains in Human HOX Loci by Noncoding RNAs. Cell 129, 1311–1323.

Rogakou, E.P., Pilch, D.R., Orr, A.H., Ivanova, V.S., and Bonner, W.M. (1998). DNA double-stranded breaks induce histone H2AX phosphorylation on serine 139. J Biol Chem 273, 5858–5868.

Saito, H., Fujita, Y., Kashida, S., Hayashi, K., and Inoue, T. (2011). Synthetic human cell fate regulation by protein-driven RNA switches. Nature communications 2, 160.

Saito, H., Kobayashi, T., Hara, T., Fujita, Y., Hayashi, K., Furushima, R., and Inoue, T. (2010). Synthetic translational regulation by an L7Ae-kink-turn RNP switch. Nature chemical biology 6, 71–78.

Sander, J.D., and Joung, J.K. (2014). CRISPR-Cas systems for editing, regulating and targeting genomes. Nat Biotechnol.

Sanjana, N.E., Cong, L., Zhou, Y., Cunniff, M.M., Feng, G., and Zhang, F. (2012). A transcription activator-like effector toolbox for genome engineering. Nat Protocols 7, 171–192.

Shoval, O., and Alon, U. (2010). SnapShot: Network Motifs. Cell 143, 326–326.e321.

Sinha, A., Hughes, K.R., Modrzynska, K.K., Otto, T.D., Pfander, C., Dickens, N.J., Religa, A.A., Bushell, E., Graham, A.L., Cameron, R., et al. (2014). A cascade of DNA-binding proteins for sexual commitment and development in Plasmodium. Nature 507, 253–257.

Smith, C.W.J., Porro, E.B., Patton, J.G., and Nadal-Ginard, B. (1989). Scanning from an independently specified branch point defines the 3[prime] splice site of mammalian introns. Nature 342, 243–247.

Soukup, G.A., and Breaker, R.R. (1999). Design of allosteric hammerhead ribozymes activated by ligand-induced structure stabilization. Structure 7, 783–791.

Sprinzak, D., Lakhanpal, A., LeBon, L., Garcia-Ojalvo, J., and Elowitz, M.B. (2011). Mutual inactivation of Notch receptors and ligands facilitates developmental patterning. PLoS computational biology 7, e1002069.

Sternberg, S.H., Haurwitz, R.E., and Doudna, J.A. (2012). Mechanism of substrate selection by a highly specific CRISPR endoribonuclease. Rna 18, 661–672.

Tabor, J.J., Salis, H.M., Simpson, Z.B., Chevalier, A.A., Levskaya, A., Marcotte, E.M., Voigt, C.A., and Ellington, A.D. (2009). A synthetic genetic edge detection program. Cell 137, 1272–1281.

Taggart, A.J., DeSimone, A.M., Shih, J.S., Filloux, M.E., and Fairbrother, W.G. (2012). Large-scale mapping of branchpoints in human pre-mRNA transcripts in vivo. Nature structural & molecular biology 19, 719–721.

Teichmann, M., Dieci, G., Pascali, C., and Boldina, G. (2010). General transcription factors and subunits of RNA polymerase III: Paralogs for promoter- and cell type-specific transcription in multicellular eukaryotes. Transcription 1, 130–135.

Upadhyay, S.K., Kumar, J., Alok, A., and Tuli, R. (2013). RNA-Guided Genome Editing for Target Gene Mutations in Wheat. G3: Genes|Genomes|Genetics 3, 2233–2238.

Urlinger, S., Baron, U., Thellmann, M., Hasan, M.T., Bujard, H., and Hillen, W. (2000). Exploring the sequence space for tetracycline-dependent transcriptional activators: Novel mutations yield expanded range and sensitivity. Proceedings of the National Academy of Sciences 97, 7963–7968.

Wang, D., Wang, H., Brown, J., Daikoku, T., Ning, W., Shi, Q., Richmond, A., Strieter, R., Dey, S.K., and DuBois, R.N. (2006). CXCL1 induced by prostaglandin E2 promotes angiogenesis in colorectal cancer. J Exp Med 203, 941–951.

Wang, H., Yang, H., Shivalila, C.S., Dawlaty, M.M., Cheng, A.W., Zhang, F., and Jaenisch, R. (2013). One-step generation of mice carrying mutations in multiple genes by CRISPR/Cas-mediated genome engineering. Cell 153, 910–918.

Wang, X., Arai, S., Song, X., Reichart, D., Du, K., Pascual, G., Tempst, P., Rosenfeld, M.G., Glass, C.K., and Kurokawa, R. (2008). Induced ncRNAs allosterically modify RNA-binding proteins in cis to inhibit transcription. Nature 454, 126–130.

Weber, W., Bacchus, W., Gruber, F., Hamberger, M., and Fussenegger, M. (2007). A novel vector platform for vitamin H-inducible transgene expression in mammalian cells. Journal of biotechnology 131, 150–158.

Weber, W., and Fussenegger, M. (2009). Engineering of Synthetic Mammalian Gene Networks. Chemistry & Biology 16, 287–297.

Weber, W., and Fussenegger, M. (2012). Emerging biomedical applications of synthetic biology. Nature reviews Genetics 13, 21–35.

Weber, W., Fux, C., Daoud-el Baba, M., Keller, B., Weber, C.C., Kramer, B.P., Heinzen, C., Aubel, D., Bailey, J.E., and Fussenegger, M. (2002). Macrolide-based transgene control in mammalian cells and mice. Nature biotechnology 20, 901–907.

Weber, W., Schoenmakers, R., Keller, B., Gitzinger, M., Grau, T., Daoud-El Baba, M., Sander, P., and Fussenegger, M. (2008). A synthetic mammalian gene circuit reveals antituberculosis compounds. Proceedings of the National Academy of Sciences of the United States of America 105, 9994–9998.

White, R.J. (1998). RNA Polymerase III Transcription (Springer-Verlag).

Willis, I.M. (1993). RNA polymerase III. European Journal of Biochemistry 212, 1–11.

Wilson, R.C., and Doudna, J.A. (2013). Molecular Mechanisms of RNA Interference. Annual Review of Biophysics 42, 217–239.

Wilusz, J.E., Freier, S.M., and Spector, D.L. (2008). 3′ End Processing of a Long Nuclear-Retained Noncoding RNA Yields a tRNA-like Cytoplasmic RNA. Cell 135, 919–932.

Wilusz, J.E., JnBaptiste, C.K., Lu, L.Y., Kuhn, C.D., Joshua-Tor, L., and Sharp, P.A. (2012). A triple helix stabilizes the 3’ ends of long noncoding RNAs that lack poly(A) tails. Genes & development 26, 2392–2407.

Win, M.N., Liang, J.C., and Smolke, C.D. (2009). Frameworks for programming biological function through RNA parts and devices. Chem Biol 16, 298–310.

Xie, Z., Wroblewska, L., Prochazka, L., Weiss, R., and Benenson, Y. (2011). Multi-input RNAi-based logic circuit for identification of specific cancer cells. Science 333, 1307–1311.

Yang, H., Wang, H., Shivalila, C.S., Cheng, A.W., Shi, L., and Jaenisch, R. (2013). One-step generation of mice carrying reporter and conditional alleles by CRISPR/Cas-mediated genome engineering. Cell 154, 1370–1379.

Ye, H., Daoud-El Baba, M., Peng, R.W., and Fussenegger, M. (2011). A synthetic optogenetic transcription device enhances blood-glucose homeostasis in mice. Science 332, 1565–1568.

Yin, Q.-F., Yang, L., Zhang, Y., Xiang, J.-F., Wu, Y.-W., Carmichael, G.G., and Chen, L.-L. (2012). Long Noncoding RNAs with snoRNA Ends. Molecular cell 48, 219–230.

Ying, S.-Y., and Lin, S.-L. (2005). Intronic microRNAs. Biochemical and Biophysical Research Communications 326, 515–520.

Zhao, J., Sun, B.K., Erwin, J.A., Song, J.-J., and Lee, J.T. (2008). Polycomb Proteins Targeted by a Short Repeat RNA to the Mouse X Chromosome. Science 322, 750–756.

